# Identification of novel glycosylation events on human serum-derived Factor IX

**DOI:** 10.1101/567941

**Authors:** Cassandra L. Pegg, Lucia F. Zacchi, Dinora Roche Recinos, Christopher B. Howard, Benjamin L. Schulz

## Abstract

Human Factor IX is a highly post-translationally modified protein that is an important clotting factor in the blood coagulation cascade. Functional deficiencies in Factor IX result in the bleeding disorder haemophilia B, which is treated with plasma-derived or recombinant Factor IX concentrates. Here, we investigated the post-translational modifications of human serum-derived Factor IX and report previously undescribed *O*-linked monosaccharide compositions at serine 141 and a novel site of glycosylation. At serine 141 we observed two monosaccharide compositions, with HexNAc_1_Hex_1_NeuAc_2_ dominant and a low level of HexNAc_1_Hex_1_NeuAc_1_. This *O*-linked site lies N-terminal to the first cleavage site for the activation peptide, an important region of the protein that is removed to activate Factor IX. The novel site is an *N*-linked site in the serine protease domain with low occupancy in a non-canonical consensus motif at asparagine 258, observed with a HexNAc_4_Hex_5_NeuAc_2_ monosaccharide composition attached. This is the first reported instance of a site of modification in the serine protease domain. The description of these glycosylation events provides a basis for future functional studies and contributes to structural characterisation of native Factor IX for the production of effective therapeutic biosimilars and biobetters.

## INTRODUCTION

Human Factor IX is a single chain zymogen that circulates in the plasma and plays an integral role in the coagulation pathway that stems bleeding at sites of vascular injury [1]. During its synthesis, Factor IX is extensively modified in the endoplasmic reticulum (ER) and Golgi apparatus including: removal of the pre-pro leader sequence, γ-carboxylation, β-hydroxylation, phosphorylation, sulfation and *N*- and *O*-linked glycosylation [2,3]. After activation of the coagulation pathway, Factor IX is converted to the active serine protease Factor IXa, which then activates Factor X to facilitate fibrin clot formation [1,4]. This activity requires Factor IX as well as the presence of Factor VIIIa. Deficiencies of functional Factor IX result in haemophilia B, a serious haemorrhagic disorder that affects approximately one in 30,000 males worldwide [5]. Current treatments for haemophilia B include infusions of serum-derived or recombinant proteins, the latter of which is the preferred treatment in high-income countries [6–9]. Disadvantages of these treatments include their high cost, the short half-life of Factor IX molecules, the potential for thrombotic episodes, and with respect to serum-derived products, the risk of viral and prion transmission [9]. Therefore, more effective recombinant versions of Factor IX are needed that overcome the drawbacks of current treatments.

A key challenge associated with producing recombinant Factor IX is the production of adequate quantities of the protein that contain the types and levels of post-translational modifications (PTMs) found on the native protein. The PTMs of Factor IX modulate the function, activity, and serum half-life of the protein [10,3]. Given the importance of PTMs and the demand for effective Factor IX biosimilars, the PTM profiles of serum-derived (Fig. 1a) and recombinant Factor IX have been extensively characterised [2,3]. Human Factor IX is synthesised as a 461 amino acid precursor molecule which is processed into a mature form through removal of the 28-residue signal peptide (SP) and 18-residue propeptide (PP). Before the PP is proteolytically removed in the *trans*-Golgi, it targets Factor IX to the vitamin K dependant enzyme γ-glutamyl-carboxylase in the ER, which catalyses carboxylation of 12 Glu residues to γ-glutamic acid (Gla) in the N-terminal Gla domain of Factor IX [11,12]. The dense cluster of Gla residues in this domain are essential for the activity of the secreted protein through coordination of Ca^2+^ ions. The protein-bound Ca^2+^ ions induce conformational changes in the Gla domain, promoting proper folding of the domain and anchoring of Factor IX to phospholipids on plasma membranes during blood coagulation [13]. Removal of the PP before secretion is needed to obtain a functionally active molecule as it enables the N-terminal Gla domain to fold into a biologically active conformation [14]. The Gla domain of Factor IX is followed by two epidermal growth factor-like (EGF) domains. Several residues are modified within the first EGF (EGF1) domain, including β-hydroxylation of Asp64 to β-hydroxyaspartate (Hya) [15,16], phosphorylation of Ser68 [17], *O*-linked glucose (Glc) elongated by xylose (Xyl) at Ser53 [18], and *O*-linked fucose (Fuc) at Ser61 [19,20]. During the coagulation process a 35-residue activation peptide (AP) C-terminal to the second EGF (EGF2) domain is cleaved from the mature protein [1]. N-terminal to the AP is a *O*-linked glycosylation site at Ser141 [21]. The AP is itself is heavily modified. It contains two sequons for *N*-linked glycosylation (Asn-Xaa-Ser/Thr; Xaa not Pro) at Asn157 and Asn167 [22] and multiple sites of *O*-linked glycosylation at Thr159 [23], Thr169 [23], and Thr172 [2]. It is noted in UniProt that Thr179 of human Factor IX (P00740) is also *O-*glycosylated, however the publication studied recombinant Factor IX produced in pigs [24] (confirmed through personal communication with the corresponding author). Sulfation (Sulf) at Tyr155 and phosphorylation (Phos) at Thr158 have also been reported [2]. Until now, PTMs of the C-terminal serine protease domain had not been described.

**Fig. 1.**
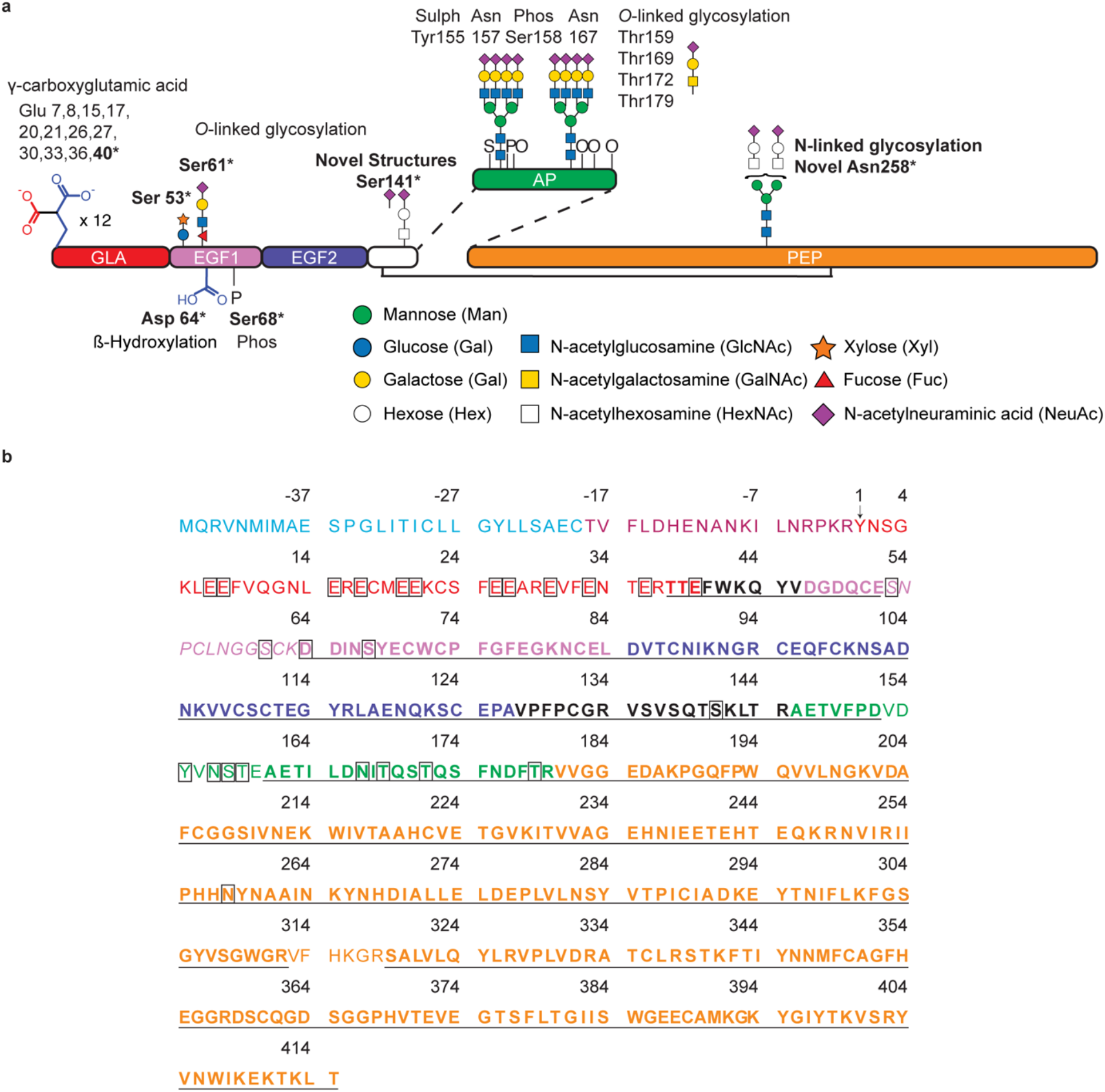
Domains, modifications, and sequence coverage of human serum-derived Factor IX. **(a)** Schematic representation of the mature form of Factor IX. The primary structure includes the N-terminal γ-carboxyglutamic acid domain (GLA), epidermal growth factor-like domains (EGF), activation peptide (AP), and serine protease domain (PEP). A disulfide bond connecting the two polypeptide chains formed after cleavage of AP is shown by a solid line. Locations of post-translational modifications are denoted with the numbering of the amino acids at the sites of attachment. Sulf, sulfation. Phos, phosphorylation. Glycans are represented following the Symbol Nomenclature for Glycans [36]. *, Modified residues that were observed in this work. (b) Amino acid sequence of Factor IX (UniProt ID P00740). The first amino acid residue of the mature form of the protein is designated residue number 1. The amino acid residues in each domain are described in [4] and are represented as follows: signal peptide (blue), propeptide (magenta), GLA domain (red) EGF1 (pink) and EGF2 (purple) domains, AP (green) and PEP domain (orange). Modified residues are highlighted in boxes. Amino acid sequence coverage of Factor IX derived from Sequest HT searches in Proteome Discoverer is shown in bold and underlined. Additional sequence coverage obtained from Byonic searches is shown in italics

With recent technical advances in mass spectrometry (MS) instrumentation, characterisation of protein PTMs including complex modifications like glycosylation have become more achievable. During protein glycosylation, oligosaccharide structures known as glycans are covalently linked to amino acid side chains [25]. Glycosylation confers additional layers of complexity to the structural and functional properties of proteins and can have a profound influence on both normal and pathological biological processes [26]. Glycans are naturally heterogeneous due to non-template driven biosynthetic processing and branching of the glycosidic linkages [27]. Both native and recombinant glycoproteins can be observed as different glycoforms with varying structures and degrees of site occupancy [28]. The two most common forms of protein glycosylation, *N*-linked and *O*-linked, stem from the conjugation of glycans to the side-chain amide nitrogen of Asn or the side chain hydroxyl group of Ser and Thr residues, respectively [27]. Unlike the *N*-linked glycosylation sequon, there is no amino acid consensus sequence that predicts likely sites for the *O*-glycosylation of Ser and Thr residues. The complex nature of protein glycosylation and subsequent structural diversity of glycoproteins can present significant challenges for characterisation of the glycosylation profile of glycoproteins [29]. Here, we used advanced MS and data analysis strategies to investigate PTMs of human serum-derived Factor IX. We confirmed *O*-linked occupancy at Ser141 and describe two new monosaccharide compositions. We also observed one novel *N*-glycosylation site, which, to our knowledge, has not been previously described.

## MATERIALS AND METHODS

### Proteolytic and endoglycosidase digestions

Human serum-derived Factor IX was purchased from Sigma Aldrich, MO, USA (Product number F0806; Batch number SLBT6630; ≥ 85% pure as determined by SDS-PAGE). This Factor IX product is purified from a concentrated pool of normal human plasma and immunopurified using a monoclonal antibody specific to Factor IX coupled to agarose. Approximately 15 μg was denatured and cysteines reduced by incubation in 6 M guanidine-HCl, 50 mM TrisHCl buffer pH 8, and 10 mM dithiothreitol with shaking for 30 min at 30 °C. The reduced proteins were alkylated by addition of acrylamide to a final concentration of 25 mM and incubation with shaking at 30 °C for 1 h. The reaction was quenched with 5 mM dithiothreitol followed by precipitation overnight at −20 °C in four volumes of methanol/acetone (1:1 v/v). The precipitated proteins were centrifuged for 10 min at 18,000 rcf and the protein pellet was resuspended in 60 μL of 50 mM NH_4_HCO_3_ and briefly vortexed. Each protein sample was split into three aliquots (5 μg) and digested with either sequencing grade porcine trypsin (Sigma-Aldrich, MO, USA), bovine chymotrypsin, or endoproteinase Glu-C from *Staphylococcus aureus* V8 (Roche Diagnostics GmbH, Mannheim, Germany). Enzyme to protein ratios were 1:20, 1:30, and 1:10, respectively, and the digests proceeded for 16 h at 37 °C. The proteolytic enzymes were inactivated by heating for 5 min at 95 °C followed by cooling to room temperature and the addition of 1 mM phenylmethylsulfonyl fluoride and incubation at room temperature for 10 min. The proteolytic digests were aliquoted into equal volumes (containing ~2.5 μg of protein each) and peptides in one aliquot were deglycosylated by further treatment with 125 U of Peptide-N-Glycosidase F (PNGase F) (New England BioLabs, MA, USA) for 16 h at 37 °C.

### Sample clean-up and mass spectrometry

Peptides were desalted and concentrated with C18 ZipTips (10 μL pipette tip with a 0.6 μL resin bed; Millipore, MA, USA). Samples were dried and reconstituted in 25 or 50 μL 0.1% formic acid depending on the specific analysis, and ~200 ng of peptides were injected for each chromatographic run using a Dionex UltiMate 3000 uHPLC system (Thermo Fisher Scientific, Bremen, Germany). Solvent A was 1% CH_3_CN in 0.1% (v/v) aqueous formic acid and solvent B was 80% (v/v) CH_3_CN containing 0.1% (v/v) formic acid. Samples were loaded onto a C18 Acclaim^TM^ PepMap^TM^ trap column (100 Å, 5 μm × 0.3 mm × 5 mm, Thermo Fisher Scientific) and washed for 3 min at 30 μL/min before peptides were eluted onto a C18 Acclaim^TM^ PepMap^TM^ column (100 Å, 5 μm × 0.75 mm × 150 mm, Thermo Fisher Scientific) at flow rate of 0.3 μL/min. Tryptic peptides were separated with a gradient of 10% to 60% solvent B over 90 min, and the remaining digests were separated with a gradient of 8% to 40% solvent B over 36 min. The samples were analysed on an Orbitrap Elite mass spectrometer (Thermo Fisher Scientific). Survey scans of peptide precursors from an *m/z* of 300 to 1800 were acquired in the Orbitrap at a resolution of 120K (full width at half-maximum, FWHM) at 400 *m/z* using an automatic gain control target of 1,000,000 and maximum injection time of 200 ms. The ten most intense precursors with charge states above two were selected for fragmentation by beam-type collision-induced dissociation (CID) (also termed higher-energy C-trap dissociation or HCD) using a normalized collision energy of 35% with a precursor isolation window of 2 Da. Fragment ions were acquired in the Orbitrap at a resolution of 30K using an automatic gain control target of 100,000 and maximum injection time of 200 ms. If the fragmented precursor ions produced diagnostic glycan oxonium ions of *N*-acetylhexosamine (HexNAc) (204.0867), HexNAc_1_-H_2_O (186.0761), Hexose (Hex) (163.0601) or HexNAc_1_Hex_1_ (366.1395) within a ±10 ppm window the precursor ions were then re-isolated and subjected to electron-transfer dissociation (ETD) with supplemental CID activation (ETciD).

### Data analysis

The Sequest HT node in Proteome Discoverer (v. 2.0.0.802 Thermo Fisher Scientific) was used to search HCD spectra from the RAW files generated by MS analysis. The protein databases used were human (UniProt UP000005640, downloaded 20 April 2018 with 20,303 reviewed proteins) with a custom contaminants database. Cleavage specificity was set as C-terminal and included trypsin (Arg/Lys except when followed by Pro), Glu-C (Asp/Glu) or chymotrypsin (Phe/Leu/Trp/Tyr). The enzyme specificity was set as semi-specific for the chymotrypsin digest, which allows non-specific cleavage at either the N- or C-terminal end of a peptide. A maximum of two missed cleavages were allowed. Mass tolerances of 10 ppm and 0.02 Da were applied to precursor and fragment ions, respectively. Cys-S-beta-propionamide was set as a static modification, and dynamic modifications were set to deamidation of Asn and mono-oxidised Met. A maximum of three dynamic modifications were allowed per peptide. Confident peptide-to-spectrum matches (PSMs) were assigned using the “Percolator” node and a maximum Delta Cn of 0.05 was applied. At least two unique peptides were required for confident protein identification. Precursor peak areas were calculated using the Precursor Ions Area Detector node, the results of which were used to calculate occupancy of post-translationally modified sites. Occupancy was calculated by the area under the curve of the total modified peptides as a percentage of the area under the curve of total modified and non-modified peptides [30].

For Byonic searches (Protein Metrics, v. 2.13.17) HCD and ETD fragmentation was selected, and cleavage specificity was set as per the Sequest HT searches. A maximum of two missed cleavages were allowed and mass tolerances of 10 ppm and 15 ppm were applied to precursor and fragment ions, respectively. Cys-S-beta-propionamide was set as a fixed modification and variable modifications were set according to the predicted number and types of modifications on a single peptide. The setting “Common 1”, which allowed each modification to be present once on a peptide, included mono-oxidised Met and Asp (Hya), phosphorylation of Ser or Thr, and sulfation of Ser, Thr, and Tyr. The setting “Common 2”, which allowed each modification to be present twice on a peptide, included deamidation of Asn and carboxy Glu. The setting “Rare 1”, which allowed each modification to be present once on a peptide, included the monosaccharide composition HexNAc_4_Hex_5_NeuAc_2_ (where NeuAc is *N*-acetylneuraminic acid) at any Asn residue and the Byonic glycan database of 57 *N*-glycans from human plasma at the consensus sequence N-X-S/T. Four additional *N*-linked structures HexNAc_6_Hex_7_NeuAc_4_, HexNAc_6_Hex_7_NeuAc_4_Fuc_1_, HexNAc_5_Hex_6_NeuAc_3_ and HexNAc_5_Hex_6_NeuAc_3_Fuc_1_ that were not in the Byonic *N-*glycan database were included as they are the most abundant glycans on Factor IX described in the literature [28,31]. A biantennary glycan with the monosaccharide composition HexNAc_4_Hex_5_NeuAc_2_ already present in the Byonic database has also been described as an abundant species from serum derived FIX [28,31]. The *O*-linked monosaccharide compositions HexNAc_1_Hex_1_, HexNAc_1_Hex_1_Fuc_1_NeuAc_1_, HexNAc_1_Hex_1_NeuAc_2_, Hex_1_Pent_1_ or Hex_1_Pent_2_, (where Pent is pentose) at any Ser or Thr residues were also allowed once per peptide. The setting “Rare 2”, which allowed each modification to be present twice on a peptide included HexNAc_1_Hex_1_NeuAc_1_ at any Ser or Thr residues. The full list of glycans used in the searches can be found in Table S1. A maximum of three common modifications and two rare modifications were allowed per peptide. The protein database contained Factor IX isoform 1 (UniProt ID P00742), isoform 2 (UniProt ID P00740-2) which has the EGF1 domain missing, Factor IX isoform 1 with the SP and PP removed, and Factor IX isoform 1 with the amino acid change 194T>A [32]. All assigned peptides with PTMs of interest were manually inspected and the best unique matches were included in the results. To ensure all glycopeptides selected for MS2 fragmentation were identified in this study validation of the Byonic results was conducted using Xcalibur Qual Browser (v. 3.0.63 Thermo Scientific). An extracted ion chromatograph for the theoretical *m/z* value of [HexNAc+H]^+^ (204.0867) was used to investigate potential glycopeptides that were not identified in Byonic searches. All MS2 spectra containing this HexNAc oxonium ion were manually inspected. For *de novo* sequencing of glycopeptides not identified in Byonic searches the theoretical masses of peptide and peptide fragment ions were calculated using the MS-Digest and MS-Product modules of Protein Prospector, respectively (http://prospector.ucsf.edu). The parameters for the Byonic searches were then changed to include previously unidentified glycopeptides resulting in the final parameters described above.

The mass spectrometry proteomics data have been deposited to the ProteomeXchange Consortium via the PRIDE [33] partner repository with the dataset identifier PXD012711.

A multiple sequence alignment was used to assess the conservation of the novel *N*-linked site from Factor IX in highly structurally homologous vitamin K-dependant proteins Factor VII, Factor X, protein Z and protein C and in Factor IX from other species (*Bos taurus*, *Pan troglodytes*, *Mus musculus*, *Canis lupus familiaris, Sus scrofa* and *Rattus norvegicus*). The sequences were retrieved from UniProt and protein sequences were aligned using Clustal Omega [34] available at http://www.uniprot.org applying the default parameters (Gonnet transition matrix, gap opening penalty of six bits and gap extension of one bit) [35].

## RESULTS AND DISCUSSION

### High sequence coverage obtained of Factor IX using proteomic searches

We used the Sequest HT node in Proteome Discoverer to assess the purity of the sample and sequence coverage of Factor IX after digestion with trypsin, Glu-C, or chymotrypsin with and without additional PNGase F *N*-glycan release. PNGase F cleaves between the innermost *N*-acetylglucosamine (GlcNAc) and Asn residues of high mannose, hybrid, and complex *N*-linked glycans, converting the previously glycosylated Asn to Asp. This cleavage event results in a mass shift of +0.984 (Asn>Asp) compared to unmodified Asn. Accordingly, Asn residues within *N*-linked consensus sites can be assigned as unmodified or modified after peptides are detected by MS. The two isoforms of Factor IX (P00740 and P00740-2) in the UniProt reference human proteome were the highest scoring proteins in all searches (Table S2). Prothrombin (UniProt ID P00734) was also identified in each of the searches, although the number of PSMs identified was considerably lower than those observed for Factor IX. For example, 148 PSMs were identified for prothrombin, compared to 2,359 PSMs for Factor IX in the trypsin digested sample (Table S-2a). Sequence coverage of Factor IX ranged between 57-67% after trypsin and Glu-C digests and 36-44% after chymotrypsin digestion (Fig. S-1). The combination of the results of the Sequest HT searches from all six digests gave 85% coverage of the mature form of the protein. The SP and PP are physiologically cleaved prior to secretion of the mature protein, and indeed PSMs from this region were not observed in any of the digests (Fig. 1b). It has been reported that there is complete modification of every Glu residue to Gla in the Gla domain of serum-derived Factor IX [2]. However, we observed one tryptic peptide from the Gla domain, T_38_TEFWK_43_, which included unmodified Glu40 (Table S-2b). Within the EGF1 domain, Ser53 and Ser61 are highly modified with *O*-linked glycans [18-20] and Asp64 undergoes partial β-hydroxylation (26-33%) [16,19]. Consistent with this, PSMs containing the two *O*-linked sites were not observed while the tryptic peptide D_64_DINSYECWPFGFEGK_80_ containing Asp64 was observed in an unmodified form. The AP contains eight reported sites of modifications, two near completely occupied [2] sites of *N*-linked glycosylation at Asn157 and Asn167, partial occupancy of *O*-linked sites Thr159 [23], Thr169 [23], Thr179 [24] and Thr172 [2] and a high degree of sulfation and/or phosphorylation [28] predicted to be at Tyr155 and Thr158 [2]. Here, we only identified PSMs from the AP after PNGase F digestion with the exception of a small unmodified region, A_146_ETVFPD_152_, at the N-terminus of the AP. Three peptide groups were observed that included various sequences from the region

A_161_ETILDN_167_IT_169_QST_172_QSFNDFT_179_RVVGGE_185_. Twelve PSMs from this region contained Asn167 and all annotated spectra for these PSMs included MS/MS fragment ions that confirmed deamidation at this site. This indicated that Asn167 was previously occupied with an *N*-linked glycan. This region of the AP also contains three predicted *O*-linked glycosylation sites at Thr69, Thr172 and Thr179. Observation of PSMs containing these sites in an unmodified form confirmed the partial occupancy described in the literature. One region from the AP was not detected, V_153_DY_155_VN_157_S_158_TEAE_162_, which contains four reported sites of modification, sulfation at Tyr155 [2], *N*-linked glycosylation at Asn157 [2], phosphorylation at Ser158 [2] and *O*-linked glycosylation at Thr159 [23]. Almost complete sequence coverage was obtained for the serine protease domain which is not predicted or reported to contain any PTMs (Fig. 1b).

### Novel glycosylation events identified on Factor IX

Using a combination of manual *de novo* sequencing and Byonic software analysis we identified previously undescribed *O*-glycan monosaccharaide compositions at Ser141 and a novel *N-*glycosylation site on Factor IX (Annotated spectra in Fig. S-2 and PSMs in Table S-3). To investigate the novel glycosylation events we used multiple proteolytic enzymes to increase coverage of the sites and two fragmentation techniques, HCD and ETciD, to confirm the monosaccharide composition of the glycans and the sites of attachment.

### *O*-glycan monosaccharide compositions and occupancy at Ser141

Occupancy at Ser141 with the monosaccharide HexNAc_1_Hex_1_ has been described in a high-throughput study of human plasma after de-sialylation and lectin enrichment of glycopeptides [21]. In this work we confirmed occupancy at Ser141 in the peptides V_135_SVSQTS_141_KLTR_145_ (after tryptic digest) and G_133_RVSVSQTS_141_KL_143_ (after chymotryptic digest) and observed two alternative monosaccharide compositions attached, HexNAc_1_Hex_1_NeuAc_2_ and HexNAc_1_Hex_1_NeuAc_1_ (Fig. 2). The site of attachment was confidently assigned to S141 through peptide sequence ions c_6_ and c_7_ after ETciD fragmentation of the tryptic glycopeptide (Fig. 2a) and the predicted monosaccharide composition of the attached glycan was supported through the observation of oxonium ions HexNAc and NeuAc (Fig. 2b). Site specificity was further confirmed through peptide sequence ions z_2_, z_3_, c_8_, and c_9_ after ETciD fragmentation of the chymotryptic glycopeptide (Fig. 2c).

**Fig. 2.**
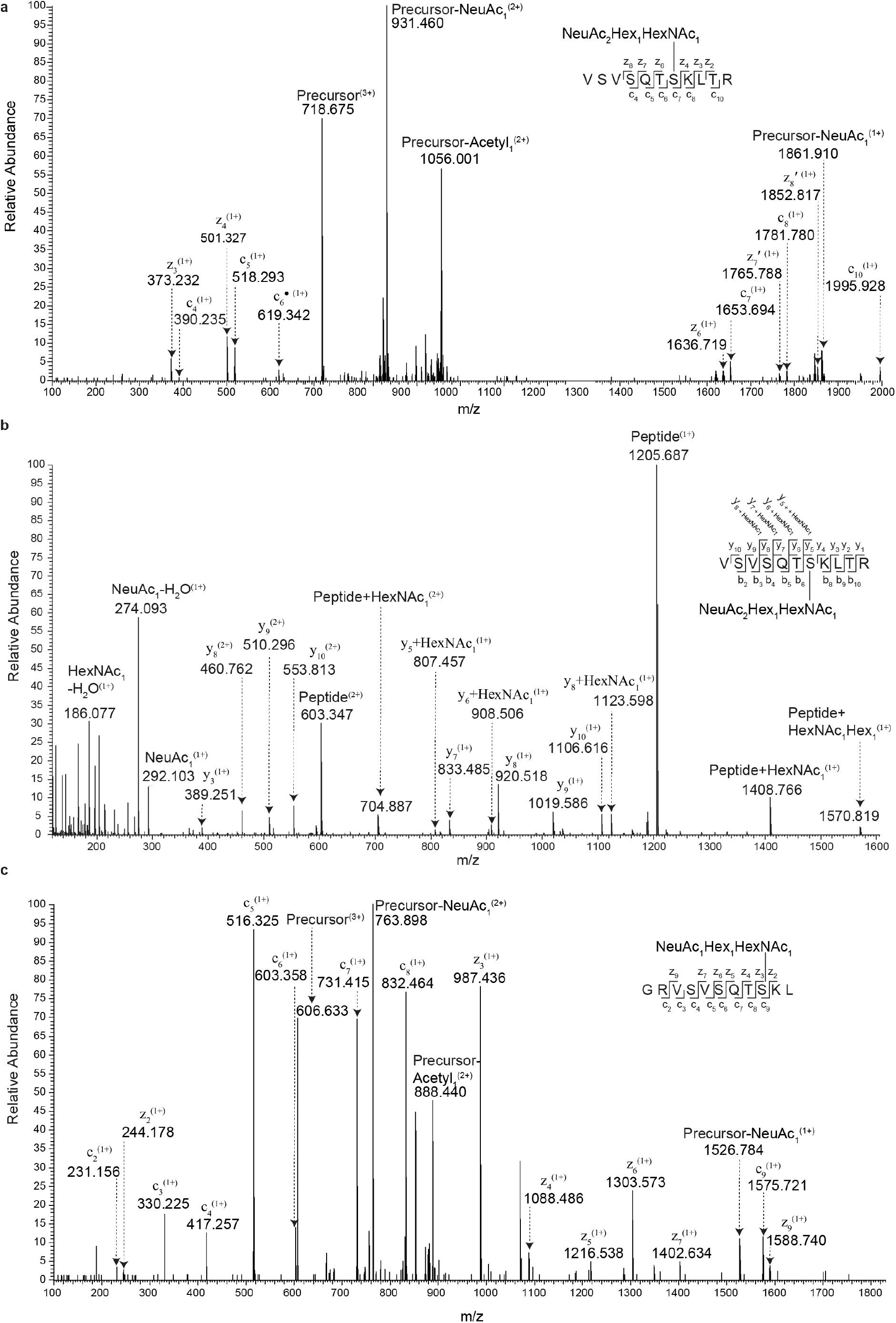
ETciD and HCD fragmentation of glycopeptides containing a the *O*-linked glycosylation site at S141 from human serum-derived Factor IX. Each panel contains a schematic of the fragmentation pattern of the glycopeptides; not all ions have been labelled in the spectra for ease of interpretation. Fragmentation of the same peptide with HexNAc_1_Hex_1_NeuAc_2_ attached at *m/z* 718.3408 (3+) with a 1.43 ppm precursor mass error using (a) ETciD and (b) HCD fragmentation. The addition of “**′**” or “•” in the labelled ETciD spectra represents even-electron z-ions or radical c-ions [38], respectively. (c) ETciD fragmentation of a glycopeptide from the chymotrypsin digest followed by PNGase F treatment with HexNAc_1_Hex_1_NeuAc_1_ attached. The precursor *m/z* value is 606.6332 (3+) with a −0.58 ppm precursor mass error

The occupancy of the *O*-linked site was approximately 83% (Table 1). The position of the *O*-linked site at S141 is intriguing as it neighbours the N-terminal region of the AP (S_141_KLTR_145_**AE**, where the first two residues of the AP are in bold). Proteolytic activation of Factor IX occurs through the removal of the 35 residue AP C-terminal to Arg145 and Arg180, producing the active serine protease designated Factor IXa [1]. The role of the *O*-glycosylation site, if any, is yet to be determined. However, it is of interest that cleavage of Factor IX by human neutrophil elastase results in a form of the protein that is not functionally active [37]. Two sites of human neutrophil elastase cleavage in Factor IX are adjacent to Ser141, at Thr140 and Thr144. *O*-linked glycosylation sites in peptides from protease-activated receptor 2 and tissue factor pathway inhibitor 1 have been shown to partially inhibit neutrophil elastase cleavage [21]. The *O*-glycosylation site in Factor IX may therefore play a role in protecting Factor IX from cleavage events that produce non-active forms of the protein.

**Table 1.** Occupancy of observed PTMs of Factor IX

| Site | Modification | Occupancy (%) |
| --- | --- | --- |
| Glu40 | γ-carboxylation | 99.3 |
| Ser53 | *O*-linked glucose | 100 |
| Ser61 | *O*-linked fucose | 100 |
| Asp64 | β-hydroxylation | 18.9 |
| Ser68 | Phosphorylation | 2.8 |
| Ser141 | *O*-linked | 83 |
|  | HexNAc<sub>1</sub>Hex<sub>1</sub>NeuAc<sub>2</sub> | 81.7 |
|  | HexNAc<sub>1</sub>Hex<sub>1</sub>NeuAc<sub>1</sub> | 1.3 |
| Asn258 | *N*-linked | 1.4 |
|  | HexNAc<sub>4</sub>Hex<sub>5</sub>NeuAc<sub>2</sub> | 1.4 |

### Novel *N*-glycosylation event identified on Factor IX

The novel *N*-linked site was observed with the monosaccharide composition HexNAc_4_Hex_5_NeuAc_2_ attached (Fig. 3). The site of attachment was assigned to Asn258 on the tryptic glycopeptide I_253_IPHHN_258_YNAAINK_265_. This peptide does not contain a *N*-linked sequon (N-X-S/T) which would easily enable site-specificity to be assumed. Nevertheless, HCD spectral evidence through peptide sequence ions b_6_+HexNAc and y_8_+HexNAc revealed the likely site of attachment was Asn258 (Fig. 3a). The production of oxonium ions for HexNAc, Hex, and NeuAc was also consistent with the proposed monosaccharide composition of the glycan (Fig. 3a). The site of attachment was more confidently assigned by ETciD where peptide sequence ions c_5_ and c_7_ and z_6-8_ confirmed that the mass of HexNAc_4_Hex_5_NeuAc_2_ was localised to Asn258 (Fig. 3b). This glycan is smaller than the most abundant *N*-glycans species observed on Factor IX in other studies [31,28,39], in which analysis of released glycans from serum-derived Factor IX found that tetrantennary and triantennary structures with varying levels of fucosylation and sialylation were abundant [31]. The occupancy of the novel *N*-linked site was approximately 1.4% (Table 1). Attachment of *N*-glycans in non-consensus motifs, although rare, has been previously documented [40], including in the heavy chain and serine protease domain of Factor XI and protein C, respectively [41,42].

**Fig. 3.**
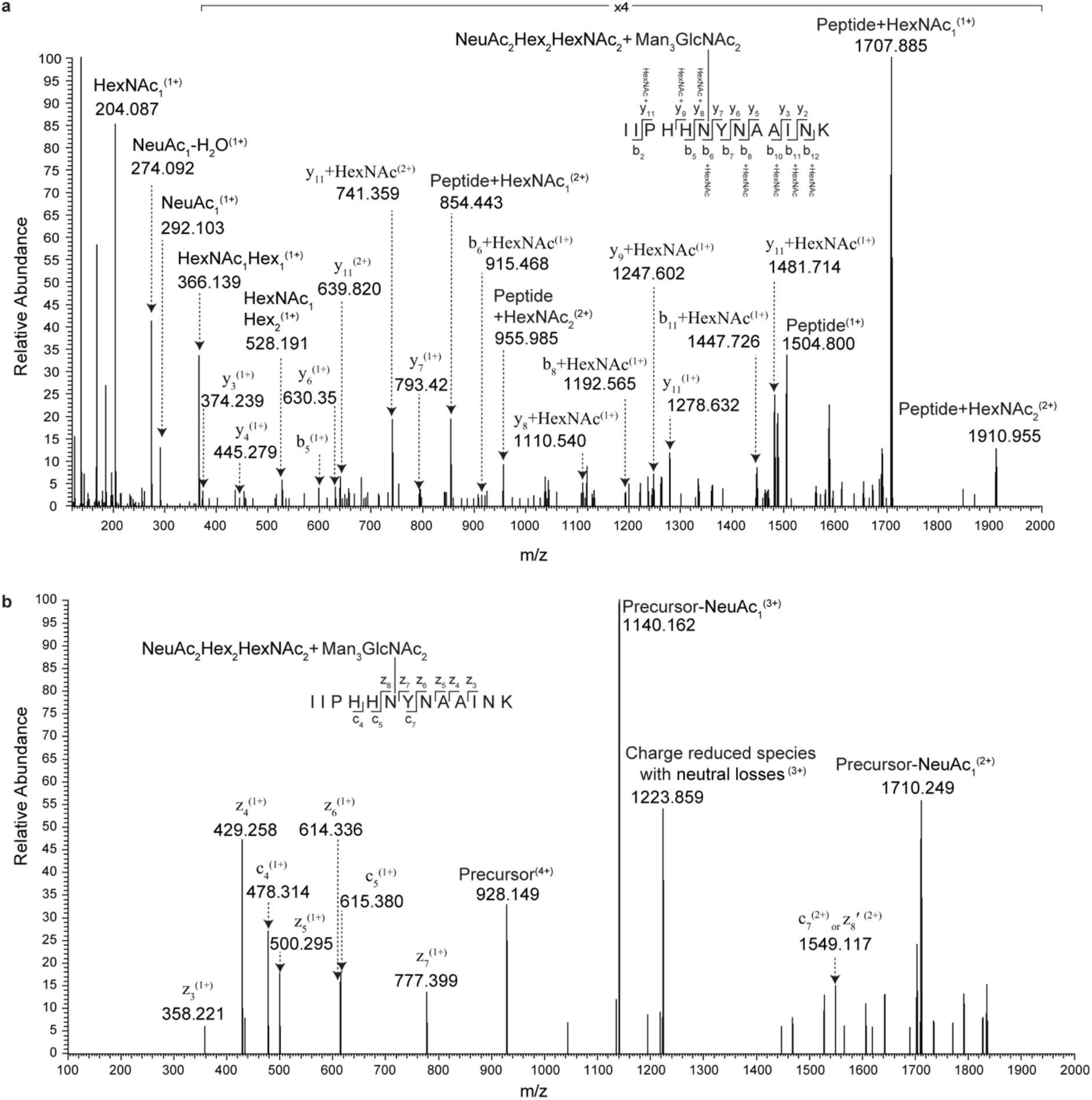
HCD and ETciD fragmentation of a glycopeptide containing a novel *N*-linked glycosylation site at Asn258 from human serum-derived Factor IX. Each panel contains a schematic of the fragmentation pattern of the glycopeptides; not all ions have been labelled in the spectra for ease of interpretation. (a) HCD fragmentation of a glycopeptide derived from the trypsin digest with HexNA_4_Hex_5_NeuAc_2_ attached. The precursor *m/z* value is 1237.1998 (3+) with a 2.80 ppm precursor mass error. (b) ETciD fragmentation of the same glycopeptide in a different charge state (4+) with a *m/z* value of 928.1516 and a 2.75 ppm precursor mass error. The addition of “**′**” in the annotated ETciD spectra represents even-electron z-ions [38]

A multiple sequence alignment was conducted to assess conservation of the novel *N*-linked site we identified in Factor IX in other human proteins that share the same domain architecture (SP-PP-Gla-EGF1-EGF2-AP-protease domain) [14] and in Factor IX from other species (Fig. S-3). We observed a conserved Asn residue at position 290 in protein C which is located within an *N*-linked consensus sequon (Fig. 4a). In addition, highly conserved *N*-linked sequons were also observed in Factor IX from other species two residues C-terminal to the novel site in human Factor IX (Fig. 4a). The Asn in protein C (position 248 in the mature form) is reported to be occupied [43] and mutation studies predict partial glycosylation [44]. Mutation of this Asn to remove the glycosylation site (N248Q) in protein C did not affect secretion of the protein but did alter the conformation of the fully processed mature protein, in particular the conformation of the active site [44]. Interestingly, the N248Q mutation in protein C also decreased intracellular cleavage of a dipeptide directly N-terminal to the AP, suggesting the *N*-linked site may alter the conformation of the unprocessed form of the protein as well. The conservation of *N*-linked consensus sites in Factor IX from other species and the observation of the partially glycosylated Asn residue in human Factor IX indicated that this region of the protein has a propensity to be *N*-glycosylated and tolerates the modification. Inspection of the crystal structure of porcine Factor IX showed that the conserved *N*-linked site sits adjacent to the active site in the serine protease domain (Fig. 4b). In Factor IX the novel *N*-linked and conserved *N*-linked sites are positioned within a surface loop (coloured magenta in Fig. 4b) that is thought to block the active site and restrict Factor IX activity in the absence of its cofactor FVIIIa [45,46]. Homology modelling using the domains of human Factor IXa revealed a similar architecture (Fig. S4). Three-dimensional modelling of the crystal structure of porcine Factor IX [47] complexed with its activator, Factor VIIa, and tissue factor [48], predicts that the catalytic site faces away from the complex interface during Factor IX activation. Thus, amino acid side chains or *N*-glycans in the surface loop may serve to restrict the activity of Factor IX during activation of the molecule. Given that glycoengineering sites into proteins can increase their half-life [49] and that the conserved *N*-linked sequon in this region of Factor IX is not deleterious in other species, it may be beneficial to introduce a similar *N*-linked sequon into recombinant forms of human Factor IX for therapeutic use.

**Fig. 4.**
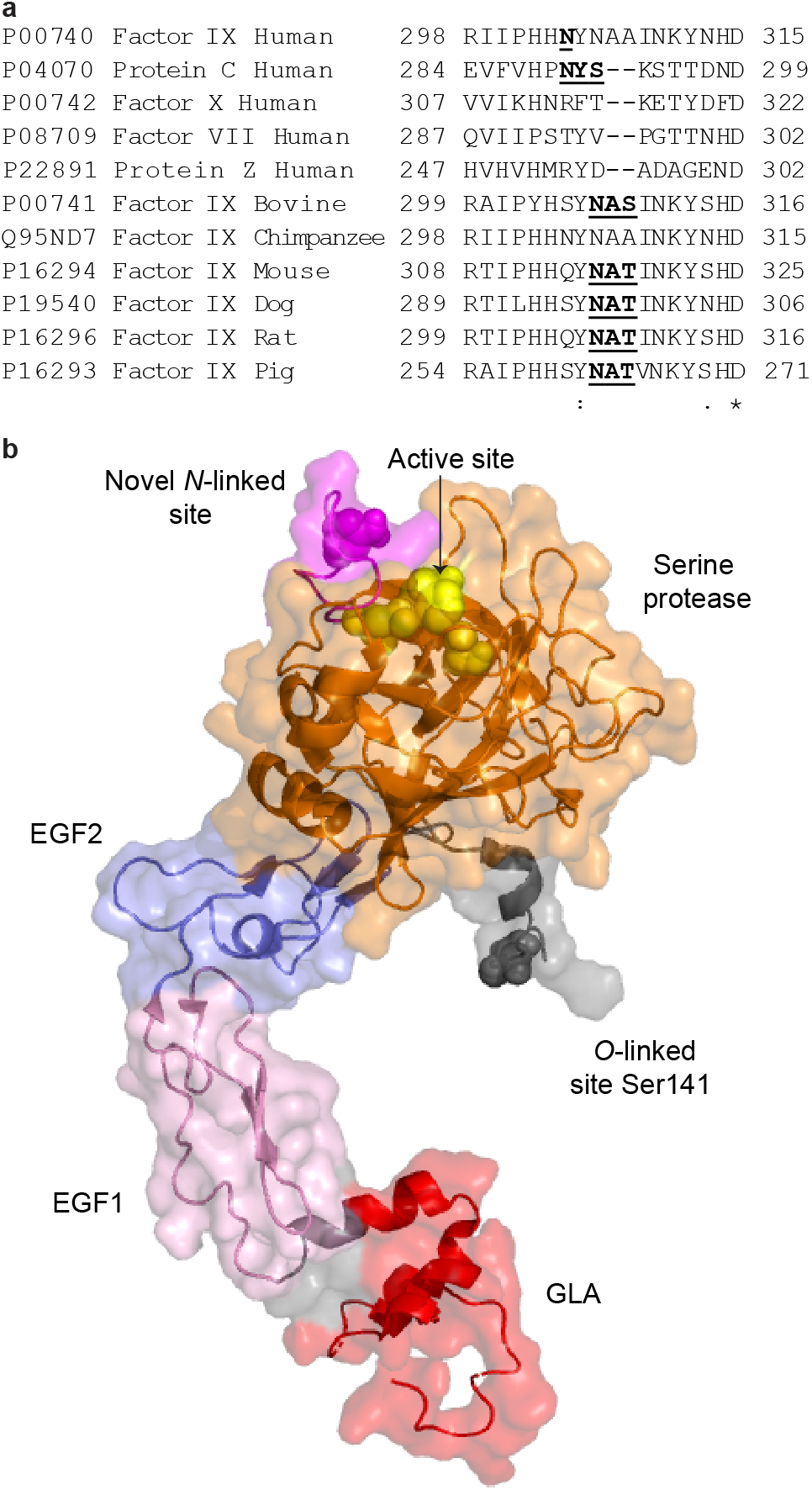
Conservation of the novel *N*-linked glycosylation site at Asn258 in human serum-derived Factor IX. (a) Multiple sequence alignment of human Factor IX homologs protein C, Factor X, Factor VII, and protein Z, as well as Factor IX from *Bos taurus* (bovine), *Pan troglodytes* (chimpanzee), *Mus musculus* (mouse), *Canis lupus familiaris* (dog), *Rattus norvegicus* (rat), and *Sus scrofa* (pig). UniProt accession numbers are listed for each protein and amino acid numbering is based on the full protein sequences. The novel *N*-linked site in human Factor IX and *N*-linked consensus sequons in the remaining proteins are in bold and underlined. (b) Crystal structure of porcine Factor IX. Surface representation of Factor IX with cartoon representation of the protein backbone (RCSB PDB identifier 1PFX [47]). The colouring of the domains follows that of Fig. 1. The GLA domain (red) anchors onto phospholipid membranes and the EGF1 (pink) and EGF2 (purple) domains form a stalk. The activation peptide is not present in this structure. The region N-terminal to the activation peptide contains a residue in the same position (grey spheres) as the *O*-linked Ser141 we observed in human serum-derived Factor IX. The serine protease domain (orange) contains the active site (yellow spheres) and a surface loop (magenta) within the protease domain includes an Asn residue (magenta spheres) within a highly conserved *N*-glycosylation sequon

### Identification of reported PTMs of Factor IX

We next used Byonic software to investigate the previously reported PTMs of Factor IX: γ-carboxylation of the first 12 Glu residues [11,12]; β-hydroxylation of Asp64 [15,16]; phosphorylation of S68 [17], *N*-linked glycosylation at Asn157 and Asn167; *O*-linked glycosylation at Ser53 [18], Ser61 [19,20], Thr159 [23], Thr169 [23], Thr179 [24], and Thr172 [2]; sulfation at Tyr155; and phosphorylation at Thr158 [2]. To accommodate the increase in search space due the numerous PTMs, we used a small protein database including only Factor IX isoforms. We judged use of this small database to be appropriate based on our confirmation of the purity of the sample by proteomic searches (Table S-2). Due the complex PTM profile of Factor IX, the parameters for the Byonic searches included eight dynamic modifications with databases of 61 *N*-linked glycans and six *O*-linked glycans. Up to five modifications were allowed per peptide and limitations were placed for specific modifications with a maximum of two of the same modification allowed per peptide. The types of variable modifications and the number of modifications allowed per peptide for the Byonic searches were selected based on the PTM profile published for Factor IX and the novel glycosylation events that we identified. All assignments were manually validated and annotated spectra are shown in Fig. S-5 and PSMs in Table S-3.

The first domain of mature serum-derived Factor IX is densely modified with Gla residues [2]. We did not characterise this Gla domain thoroughly here, even after inclusion of γ-carboxylation in the Byonic searches. It has been established that peptides containing Gla residues have decreased ionisation efficiency compared to those containing unmodified Glu residues during positive mode ESI-MS [50]. This is likely due to the negative charge imparted by the Gla residues, which has also been shown to hinder proteolytic cleavage [50], again decreasing the likelihood of detecting peptides from this region. The only modified Glu residue we identified in the Gla domain was Glu40 positioned in the tryptic peptide TTE_40_FWK (Fig. S-5A) with 99.3% occupancy (Table 1). This level of occupancy is consistent with other work reporting complete modification of every Glu residue in the Gla domain [2,51]. It has been shown that recombinant Factor IX lacking γ-carboxylation at Glu40, or at both Glu36 and Glu40, retains coagulation activity and the ability to activate factor × [51]. The tryptic peptide observed is the only theoretical peptide from the Gla region to contain a single Glu residue, and thus the least likely to suffer from the effects of decreased ionisation efficiency due to the Gla modification.

Predicted tryptic peptides from the Gla domain contain up to three possible Gla residues. To ensure our failure to identify the modified version of these peptides was not due to an artefact of search parameters, we also performed Byonic searches where up to three Gla residues were allowed per peptide. However, this also failed to identify Gla-modified peptides from this region.

The EGF1 domain of serum-derived Factor IX contains three well documented sites of modification. Two *O*-linked sites at Ser53 and Ser61 are highly occupied with the glycans Xyl_1_Glc_1_ [18] and NeuAc_1_Gal_1_GlcNAc_1_Fuc_1_ [19,20] (where Gal is galactose), respectively. We did not observe these sites in an unmodified form, suggesting they are modified with full occupancy (Table 1). The third well-documented site of modification at Asp64 undergoes 26-33% β-hydroxylation [16,19]. Using Byonic, we observed β-hydroxylation of Asp in the tryptic peptide D_64_DINSYECWPFGFEGK_80_ (Fig. S-5B) with 18.9% occupancy (Table 1). However, we were unable to unambiguously assign site-specificity between Asp64 and Asp65. The peptide S_53_NPCLNGGS_61_CKD_64_DINSYE_70_ derived from the Glu-C digest was observed with and without β-hydroxylation of Asp (Fig. S-5C and S-5D). Both observed versions of this peptide, which contain *O*-linked sites Ser53 and Ser61 along with the site of β-hydroxylation at Asp64, were additionally modified with predicted monosaccharide compositions Xyl_1_Hex_1_ and NeuAc_1_Hex_1_HexNAc_1_Fuc_1_ (we assume all deoxyhexose is fucose, and all pentose is xylose). The HCD spectrum in Fig. 5 identified the β-hydroxylated version of the glycopeptide, where the precursor mass indicated the following modifications were present: Xyl_1_Hex_1_ (294.095 Da), NeuAc_1_Hex_1_HexNAc_1_Fuc_1_ (802.286 Da), β-hydroxyaspartate (15.995 Da), and deamidation of Asn (0.984 Da). Glycan oxonium ions confirmed the presence of HexNAc, Hex, and NeuAc, but not Xyl or Fuc. The site of deamidation could be assigned to Asn58 and β-hydroxylation to either Asp64 or Asp65. The sites of *O*-linked modifications could not be determined from the fragmentation pattern. The precursor *m/z* value for this glycopeptide exceeded 850 and was doubly protonated, characteristics that have been shown to limit the effectiveness of ETD [52]. A chymotryptic glycopeptide with the peptide sequence G_59_GS_61_CKDDINSY_69_ was also observed using HCD with NeuAc_1_Hex_1_HexNAc_1_Fuc_1_ attached (Fig. S-5E). This supported previous reports [19,20] that Ser61 is the site of attachment for the glycan NeuAc_1_Gal_1_GlcNAc_1_Fuc_1_, although we could not rule out modification at Ser68. Through a process of elimination, we also confirmed that Ser53 was the site of attachment for the predicted structure Xyl_1_Glc_1_.

**Fig. 5.**
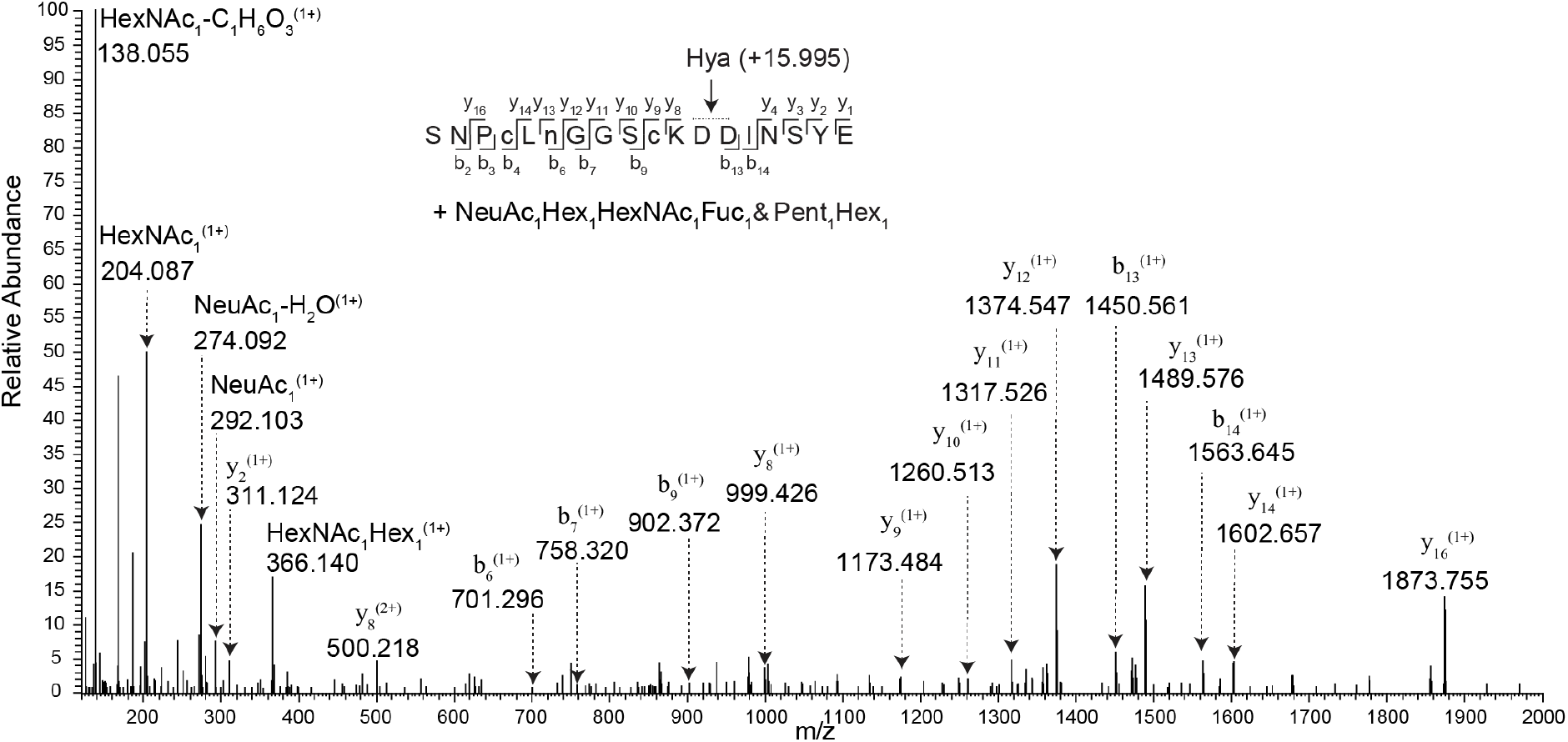
HCD fragmentation of a glycopeptide from human serum-derived Factor IX containing *O*-linked sites Ser53 and Ser61 and the site of β-hydroxylation at Asp64. The panel contains a schematic of the fragmentation pattern of the glycopeptide. c, cys-S-beta-propionamide. n, deamidation of Asn. Not all ions have been labelled in the spectra for ease of interpretation. The precursor *m/z* value is 1586.11772 (2+) with a 4.35 ppm precursor mass error

Phosphorylation of Ser68 in the EGF1 domain has also been reported [17]. The Byonic searches we conducted identified sulfation at Tyr69 in the EGF1 domain on the peptide D_64_DINS_68_Y_69_ECWPFGFEGK_80_ (Fig. S-5F) with 2.8% occupancy (Table 1, annotated as phosphorylation to align with the literature). The distinction between phosphorylation or sulfation could not be made by precursor mass alone due to the isobaric masses of the modifications at the resolution of our analyses. Furthermore, as the modified peptide was fragmented by HCD site-specificity could not be determined between Ser68 and Tyr69.

The AP is a heavily modified polypeptide. A previous compositional analysis of the amino acid and carbohydrate content revealed approximately half the mass of the AP is attributed to Hex, HexNAc, and NeuAc [1]. The major contribution of monosaccharides is likely from *N*-linked glycans attached at both Asn157 and Asn167. Released *N*-glycans of human Factor IX have been structurally and quantitatively profiled, revealing the most abundant forms to be large sialylated tetranntenary and trianntenary glycans [28,31]. Despite using three different proteases to increase coverage of the AP we were unable to detect any glycopeptides containing sites Asn157 and Asn167. As reported in the proteomic section, we only observed site Asn167 in a deamidated form and only after PNGase F digestion, indicating the site was previously occupied with an *N*-linked glycan. The lack of detection of the site in a modified form may be due to the large glycan structures and the close proximity of several other modifications including *O*-linked glycosylation. The *O*-glycan structure NeuAc_1_Gal_1_GalNAc_1_ can be attached at Thr159 and Thr169 [23] with an additional structure, Gal_1_GalNAc_1_, attached at Thr159. For ID P00740 in UniProt it is also reported that Thr179 is modified with the *O-*glycan monosaccharide compositions HexNAc_1_Hex_1_NeuAc_2_ and HexNAc_1_Hex_1_NeuAc_1_ on recombinant Factor IX produced in pigs [24]. To our knowledge, this is yet to be confirmed in plasma-derived human Factor IX. *O*-linked glycosylation at Thr172 has also been reported [2]. We observed unmodified peptides containing *O*-linked sites Thr169, Thr172 and Thr179 which indicates partial occupancy at all three sites. Sulfation at Tyr155 and phosphorylation at Thr158 have also been described [2] and using MALDI-TOF MS, an estimated 70% of total deglycosylated AP from human Factor IX was found to be modified by two phosphorylation/sulfation events [28]. A further 30% was estimated to be modified by a single phosphorylation/sulfation event. We allowed dual phosphorylation/sulfation modifications along with deamidated Asn and *O*-linked glycosylation on a single peptide in our Byonic searches. However, none of the aforementioned modifications of the AP were identified. The lack of identified PTMs from the AP may be due to the structural complexity of the region and the different methods we used when compared to other published targeted analyses such as the intact protein analytical approach for sulfation and phosphorylation of the AP [28] and amino acid and monosaccharide compositional analyses combined with fast atom bombardment MS for the characterisation of *O*-linked glycosylation at Thr159 and Thr169 [23].

## CONCLUSIONS

Our results emphasize the complexity of post-translational modifications of Factor IX. The identification of novel *O*-linked monosaccharide compositions at Ser141 and the description of a novel *N*-linked site will provide the basis for further research on the role glycosylation plays in the biological functions and regulation of Factor IX activity, and will enable comparisons between human serum-derived Factor IX and therapeutic biosimilars.

## Supporting information

ESM_1 Tables S1, S2 and S3 Byonic Glycan database, Sequest HT and Byonic Results

ESM_2 Supplementary Figures S1-S5

## NOTES

## Author Contribution

BLS, CLP, CBH, and LFZ conceived and coordinated this study. All authors designed experiments. CLP performed the experiments and analysed the data. BLS, CLP, LFZ, and DRR reviewed and interpreted the results. CLP and BLS wrote the manuscript. All authors reviewed the results and approved the final version of the manuscript. The authors declare that they have no conflicts of interest with the contents of this article.

## Acknowledgments

We thank Dr Amanda Nouwens and Peter Josh at The University of Queensland, School of Chemistry and Molecular Biosciences Mass Spectrometry Facility for their assistance and expertise.

## Supplementary material

ESM_1 Tables S1, S2 and S3 Byonic Glycan database, Sequest HT and Byonic Results

ESM_2 Supplementary Figures S1-S5

## Abbreviations

AP: activation peptide
CID: collision-induced dissociation
EGF: epidermal growth factor-like
ER: endoplasmic reticulum
ETD: electron-transfer dissociation
Fuc: fucose
Gal: galactose
Gla: γ-glutamic acid
Glc: glucose
GlcNAc: N-acetylglucosamine
HCD: higher-energy C-trap dissociation or beam-type collision-induced dissociation
Hex: hexose
HexNAc: N-acetyl-hexosamine
Hya: β-hydroxyaspartate
NeuAc: N-acetylneuraminic acid
Pent: pentose
Phos: phosphorylation
PNGase F: Peptide-N-Glycosidase F
PP: propeptide
PTMs: post-translantional modifications
PSMs: peptide-to-spectrum matches
SP: signal peptide
Sulf: sulfation
Xyl: xylose

## REFERENCES

1. Di Scipio, R.G., Kurachi, K., Davie, E.W.: Activation of human factor IX (Christmas factor). J. Clin. Invest. 61(6), 1528–1538 (1978). doi:10.1172/JCI109073

2. Bond, M., Jankowski, M., Patel, H., Karnik, S., Strang, A., Xu, B., Rouse, J., Koza, S., Letwin, B., Steckert, J., Amphlett, G., Scoble, H.: Biochemical characterization of recombinant factor IX. Semin. Hematol. 35(2 Suppl 2), 11–17 (1998).

3. Arruda, V.R., Hagstrom, J.N., Deitch, J., Heiman-Patterson, T., Camire, R.M., Chu, K., Fields, P.A., Herzog, R.W., Couto, L.B., Larson, P.J., High, K.A.: Posttranslational modifications of recombinant myotube-synthesized human factor IX. Blood 97(1), 130–138 (2001). doi:10.1182/blood.V97.1.130

4. Mathur, A., Zhong, D., Sabharwal, A.K., Smith, K.J., Bajaj, S.P.: Interaction of factor IXa with factor VIIIa. Effects of protease domain Ca2+ binding site, proteolysis in the autolysis loop, phospholipid, and factor X. J. Biol. Chem. 272(37), 23418–23426 (1997). doi:10.1074/jbc.272.37.23418

5. Goodeve, A.C.: Hemophilia B: molecular pathogenesis and mutation analysis. J. Thromb. Haemost. 13(7), 1184–1195 (2015). doi:10.1111/jth.12958

6. Lyseng-Williamson, K.A.: Coagulation factor IX (Recombinant), albumin fusion protein (Albutrepenonacog Alfa; Idelvion®): A review of its use in haemophilia B. Drugs 77(1), 97–106 (2016). doi:10.1007/s40265-016-0679-8

7. Turecek, P.L., Abbuhl, B., Tangada, S.D., Chapman, M., Gritsch, H., Rottensteiner, H., Schrenk, G., Mitterer, A., Dietrich, B., Hollriegl, W., Schiviz, A., Horling, F., Reipert, B.M., Muchitsch, E.M., Pavlova, B.G., Scheiflinger, F.: Nonacog gamma, a novel recombinant factor IX with low factor IXa content for treatment and prophylaxis of bleeding episodes. Expert Rev. Clin. Pharmacol. 8(2), 163–177 (2015). doi:10.1586/17512433.2015.1011126

8. Cooley, B., Funkhouser, W., Monroe, D., Ezzell, A., Mann, D.M., Lin, F.C., Monahan, P.E., Stafford, D.W.: Prophylactic efficacy of BeneFIX vs Alprolix in hemophilia B mice. Blood 128(2), 286–292 (2016). doi:10.1182/blood-2016-01-696104

9. Franchini, M.: Current management of hemophilia B: recommendations, complications and emerging issues. Expert Rev. Hematol. 7(5), 573–581 (2014). doi:10.1586/17474086.2014.947955

10. Johansson, L., Karpf, D.M., Hansen, L., Pelzer, H., Persson, E.: Activation peptides prolong the murine plasma half-life of human factor VII. Blood 117(12), 3445–3452 (2011). doi:10.1182/blood-2010-06-290098

11. Stanley, T.B., Wu, S.-M., Houben, R.J.T.J., Mutucumarana, V.P., Stafford, D.W.: Role of the propeptide and γ-glutamic acid domain of factor ix for in vitro carboxylation by the vitamin k-dependent carboxylase. Biochemistry 37(38), 13262–13268 (1998). doi:10.1021/bi981031y

12. Jorgensen, M.J., Cantor, A.B., Furie, B.C., Brown, C.L., Shoemaker, C.B., Furie, B.: Recognition site directing vitamin K-dependent gamma-carboxylation resides on the propeptide of factor IX. Cell 48(2), 185–191 (1987). doi:org/10.1016/0092-8674(87)90422-3

13. Huang, M., Rigby, A.C., Morelli, X., Grant, M.A., Huang, G., Furie, B., Seaton, B., Furie, B.C.: Structural basis of membrane binding by Gla domains of vitamin K-dependent proteins. Nat. Struct. Biol. 10(9), 751–756 (2003). doi:10.1038/nsb971

14. Hansson, K., Stenflo, J.: Post-translational modifications in proteins involved in blood coagulation. J. Thromb. Haemost. 3(12), 2633–2648 (2005). doi:10.1111/j.1538-7836.2005.01478.x

15. Derian, C.K., VanDusen, W., Przysiecki, C.T., Walsh, P.N., Berkner, K.L., Kaufman, R.J., Friedman, P.A.: Inhibitors of 2-ketoglutarate-dependent dioxygenases block aspartyl beta-hydroxylation of recombinant human factor IX in several mammalian expression systems. J. Biol. Chem. 264(12), 6615–6618 (1989).

16. Fernlund, P., Stenflo, J.: Beta-hydroxyaspartic acid in vitamin K-dependent proteins. J. Biol. Chem. 258(20), 12509–12512 (1983).

17. Harris R.J., P.D.I., Truong L., Smith K.J.: Partial phosphorylation of serine-68 in EGF-1 of human factor IX. In: Proceedings of XIth international conference on methods in protein structure analysis 1996

18. Nishimura, H., Kawabata, S., Kisiel, W., Hase, S., Ikenaka, T., Takao, T., Shimonishi, Y., Iwanaga, S.: Identification of a disaccharide (Xyl-Glc) and a trisaccharide (Xyl2-Glc) O-glycosidically linked to a serine residue in the first epidermal growth factor-like domain of human factors VII and IX and protein Z and bovine protein Z. J. Biol. Chem. 264(34), 20320–20325 (1989).

19. Harris, R.J., van Halbeek, H., Glushka, J., Basa, L.J., Ling, V.T., Smith, K.J., Spellman, M.W.: Identification and structural analysis of the tetrasaccharide NeuAcα (2→ 6) Galβ (1→ 4) GlcNAcβ (1→ 3) Fucα1→ O-linked to serine 61 of human factor IX. Biochemistry 32(26), 6539–6547 (1993). doi:10.1021/bi00077a007

20. Nishimura, H., Takao, T., Hase, S., Shimonishi, Y., Iwanaga, S.: Human factor IX has a tetrasaccharide O-glycosidically linked to serine 61 through the fucose residue. J. Biol. Chem. 267(25), 17520–17525 (1992).

21. King, S.L., Joshi, H.J., Schjoldager, K.T., Halim, A., Madsen, T.D., Dziegiel, M.H., Woetmann, A., Vakhrushev, S.Y., Wandall, H.H.: Characterizing the O-glycosylation landscape of human plasma, platelets, and endothelial cells. Blood Adv 1(7), 429–442 (2017). doi:10.1182/bloodadvances.2016002121

22. Kurachi, K., Davie, E.W.: Isolation and characterization of a cDNA coding for human factor IX. Proc. Natl. Acad. Sci. U. S. A. 79(21), 6461–6464 (1982).

23. Agarwala, K.L., Kawabata, S., Takao, T., Murata, H., Shimonishi, Y., Nishimura, H., Iwanaga, S.: Activation peptide of human factor IX has oligosaccharides O-glycosidically linked to threonine residues at 159 and 169. Biochemistry 33(17), 5167–5171 (1994). doi:10.1021/bi00183a021

24. Huang, L.J., Lin, J.H., Tsai, J.H., Chu, Y.Y., Chen, Y.W., Chen, S.L., Chen, S.H.: Identification of protein O-glycosylation site and corresponding glycans using liquid chromatography-tandem mass spectrometry via mapping accurate mass and retention time shift. J. Chromatogr. A 1371(Supplement C), 136–145 (2014). doi:10.1016/j.chroma.2014.10.046

25. Moremen, K.W., Tiemeyer, M., Nairn, A.V.: Vertebrate protein glycosylation: diversity, synthesis and function. Nat. Rev. Mol. Cell Biol. 13(7), 448–462 (2012). doi:10.1038/nrm3383

26. Thaysen-Andersen, M., Packer, N.H., Schulz, B.L.: Maturing glycoproteomics technologies provide unique structural insights into the N-glycoproteome and its regulation in health and disease. Mol. Cell. Proteomics 15(6), 1773–1790 (2016). doi:10.1074/mcp.O115.057638

27. Bertozzi, C.R., Rabuka, D.: Structural basis of glycan diversity. In: Varki, A., Cummings, R.D., Esko, J.D., Freeze, H.H., Stanley, P., Bertozzi, C.R., Hart, G.W., Etzler, M.E. (eds.) Essentials of Glycobiology. Cold Spring Harbor Laboratory Press, New York, NY, (2009)

28. Chevreux, G., Faid, V., Andre, M.H., Tellier, Z., Bihoreau, N.: Differential investigations from plasma-derived and recombinant Factor IX revealed major differences in post-translational modifications of activation peptides. Vox Sang. 104(2), 171–174 (2013). doi:10.1111/j.1423-0410.2012.01649.x

29. Lee, L.Y., Moh, E.S., Parker, B.L., Bern, M., Packer, N.H., Thaysen-Andersen, M.: Toward automated N-glycopeptide identification in glycoproteomics. J. Proteome Res. 15(10), 3904–3915 (2016). doi:10.1021/acs.jproteome.6b00438

30. Schulz, B.L., Aebi, M.: Analysis of glycosylation site occupancy reveals a role for Ost3p and Ost6p in site-specific N-glycosylation efficiency. Mol. Cell. Proteomics 8(2), 357–364 (2009). doi:10.1074/mcp.M800219-MCP200

31. Makino, Y., Omichi, K., Kuraya, N., Ogawa, H., Nishimura, H., Iwanaga, S., Hase, S.: Structural analysis of N-linked sugar chains of human blood clotting factor IX. J. Biochem. 128(2), 175–180 (2000). doi:10.1093/oxfordjournals.jbchem.a022738

32. McGraw, R.A., Davis, L.M., Noyes, C.M., Lundblad, R.L., Roberts, H.R., Graham, J.B., Stafford, D.W.: Evidence for a prevalent dimorphism in the activation peptide of human coagulation factor IX. Proc. Natl. Acad. Sci. U. S. A. 82(9), 2847–2851 (1985). doi:10.1073/pnas.82.9.2847

33. Perez-Riverol, Y., Csordas, A., Bai, J., Bernal-Llinares, M., Hewapathirana, S., Kundu, D.J., Inuganti, A., Griss, J., Mayer, G., Eisenacher, M., Perez, E., Uszkoreit, J., Pfeuffer, J., Sachsenberg, T., Yilmaz, S., Tiwary, S., Cox, J., Audain, E., Walzer, M., Jarnuczak, A.F., Ternent, T., Brazma, A., Vizcaino, J.A.: The PRIDE database and related tools and resources in 2019: improving support for quantification data. Nucleic Acids Res. 47(D1), D442–D450 (2019). doi:10.1093/nar/gky1106

34. Sievers, F., Wilm, A., Dineen, D., Gibson, T.J., Karplus, K., Li, W., Lopez, R., McWilliam, H., Remmert, M., Soding, J., Thompson, J.D., Higgins, D.G.: Fast, scalable generation of high-quality protein multiple sequence alignments using Clustal Omega. Mol. Syst. Biol. 7(1), 539 (2011). doi:10.1038/msb.2011.75

35. Soding, J.: Protein homology detection by HMM-HMM comparison. Bioinformatics 21(7), 951–960 (2005). doi:10.1093/bioinformatics/bti125

36. Varki, A., Cummings, R.D., Aebi, M., Packer, N.H., Seeberger, P.H., Esko, J.D., Stanley, P., Hart, G., Darvill, A., Kinoshita, T., Prestegard, J.J., Schnaar, R.L., Freeze, H.H., Marth, J.D., Bertozzi, C.R., Etzler, M.E., Frank, M., Vliegenthart, J.F., Lutteke, T., Perez, S., Bolton, E., Rudd, P., Paulson, J., Kanehisa, M., Toukach, P., Aoki-Kinoshita, K.F., Dell, A., Narimatsu, H., York, W., Taniguchi, N., Kornfeld, S.: Symbol nomenclature for graphical representations of glycans. Glycobiology 25(12), 1323–1324 (2015). doi:10.1093/glycob/cwv091

37. Samis, J.A., Kam, E., Nesheim, M.E., Giles, A.R.: Neutrophil elastase cleavage of human factor IX generates an activated factor IX-like product devoid of coagulant function. Blood 92(4), 1287–1296 (1998).

38. Zhurov, K.O., Fornelli, L., Wodrich, M.D., Laskay, U.A., Tsybin, Y.O.: Principles of electron capture and transfer dissociation mass spectrometry applied to peptide and protein structure analysis. Chem. Soc. Rev. 42(12), 5014–5030 (2013). doi:10.1039/c3cs35477f

39. Gil, G.C., Velander, W.H., Van Cott, K.E.: Analysis of the N-glycans of recombinant human Factor IX purified from transgenic pig milk. Glycobiology 18(7), 526–539 (2008). doi:10.1093/glycob/cwn035

40. Dutta, D., Mandal, C., Mandal, C.: Unusual glycosylation of proteins: Beyond the universal sequon and other amino acids. Biochim. Biophys. Acta 1861(12), 3096–3108 (2017). doi:10.1016/j.bbagen.2017.08.025

41. Faid, V., Denguir, N., Chapuis, V., Bihoreau, N., Chevreux, G.: Site-specific N-glycosylation analysis of human factor XI: Identification of a noncanonical NXC glycosite. Proteomics 14(21-22), 2460–2470 (2014). doi:10.1002/pmic.201400038

42. Miletich, J.P., Broze, G.J.: Beta protein C is not glycosylated at asparagine 329. The rate of translation may influence the frequency of usage at asparagine-X-cysteine sites. J. Biol. Chem. 265(19), 11397–11404 (1990).

43. Liu, T., Qian, W.J., Gritsenko, M.A., Camp, D.G., 2nd, Monroe, M.E., Moore, R.J., Smith, R.D.: Human plasma N-glycoproteome analysis by immunoaffinity subtraction, hydrazide chemistry, and mass spectrometry. J. Proteome Res. 4(6), 2070–2080 (2005). doi:10.1021/pr0502065

44. Grinnell, B.W., Walls, J.D., Gerlitz, B.: Glycosylation of human protein C affects its secretion, processing, functional activities, and activation by thrombin. J. Biol. Chem. 266(15), 9778–9785 (1991).

45. Kolkman, J.A., Mertens, K.: Insertion loop 256−268 in coagulation factor IX restricts enzymatic activity in the absence but not in the presence of factor VIII. Biochemistry 39(25), 7398–7405 (2000). doi:10.1021/bi992735q

46. Ponder, K.P.: FIXing factor VIII inhibitors. Blood 119(2), 325–326 (2012). doi:10.1182/blood-2011-11-389486

47. Brandstetter, H., Bauer, M., Huber, R., Lollar, P., Bode, W.: X-ray structure of clotting factor IXa: active site and module structure related to Xase activity and hemophilia B. Proceedings of the National Academy of Sciences 92(21), 9796–9800 (1995). doi:10.1073/pnas.92.21.9796

48. Chen, S.W.W., Pellequer, J.L., Schved, J.F., Giansily-Blaizot, M.: Model of a ternary complex between activated factor VII, tissue factor and factor IX. Thromb. Haemost. 88(1), 74–82 (2002).

49. Elliott, S., Lorenzini, T., Asher, S., Aoki, K., Brankow, D., Buck, L., Busse, L., Chang, D., Fuller, J., Grant, J., Hernday, N., Hokum, M., Hu, S., Knudten, A., Levin, N., Komorowski, R., Martin, F., Navarro, R., Osslund, T., Rogers, G., Rogers, N., Trail, G., Egrie, J.: Enhancement of therapeutic protein in vivo activities through glycoengineering. Nat. Biotechnol. 21(4), 414–421 (2003). doi:10.1038/nbt799

50. Hallgren, K.W., Zhang, D., Kinter, M., Willard, B., Berkner, K.L.: Methylation of gamma-carboxylated Glu (Gla) allows detection by liquid chromatography-mass spectrometry and the identification of Gla residues in the gamma-glutamyl carboxylase. J. Proteome Res. 12(6), 2365–2374 (2013). doi:10.1021/pr3003722

51. Gillis, S., Furie, B.C., Furie, B., Patel, H., Huberty, M.C., Switzer, M., Foster, W.B., Scoble, H.A., Bond, M.D.: Gamma-carboxyglutamic acids 36 and 40 do not contribute to human factor IX function. Protein Sci. 6(1), 185–196 (1997). doi:10.1002/pro.5560060121

52. Good, D.M., Wirtala, M., McAlister, G.C., Coon, J.J.: Performance characteristics of electron transfer dissociation mass spectrometry. Mol. Cell. Proteomics 6(11), 1942–1951 (2007). doi:10.1074/mcp.M700073-MCP200

