## Supplementary material for "Identification of novel glycosylation events on human serum-derived Factor IX": ESM_2 Supplementary Figures S1-S5

Glycoconjugate Journal

[†]School of Chemistry and Molecular Biosciences, The University of Queensland, St Lucia, QLD 4072, Australia.

[‡]Centre for Biopharmaceutical Innovation, Australian Institute for Bioengineering and Nanotechnology, The University of Queensland, St Lucia, QLD 4072, Australia.

Cassandra L. Pegg,[†] Lucia F. Zacchi,[‡] Dinora Roche Recinos[‡] Christopher B. Howard,[‡] and Benjamin L. Schulz[†‡*]

**Figure S-1** Protein sequence coverage of serum derived human Factor IX. Identified peptides are from HCD MS/MS analyses of samples digested with trypsin (A), trypsin followed by PNGase F (B), Glu-C (C), Glu-C followed by PNGase F (D), chymotrypsin (E) and chymotrypsin followed by PNGase F (F). Sequence coverage derived from each Sequest HT search is highlighted in green. For chymotrypsin searches with and without PNGase F cleavage specificity was set as "semi-tryptic".

(A) Trypsin

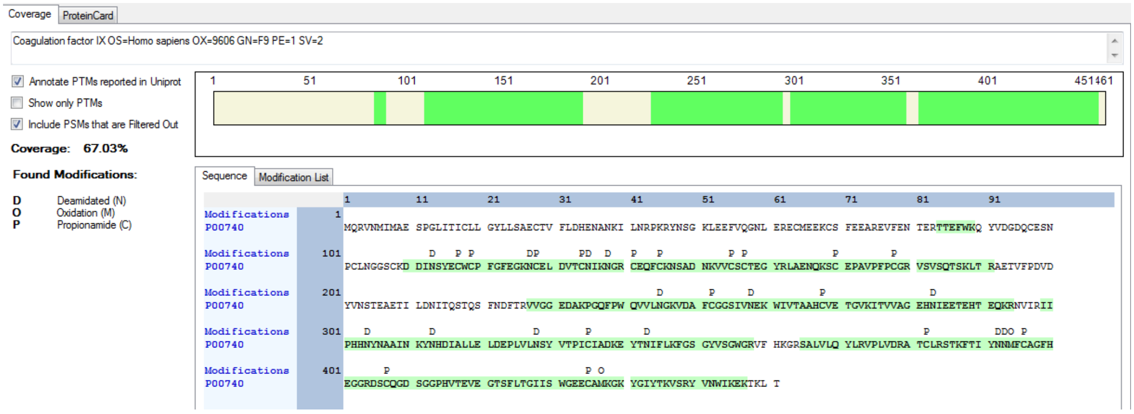


(B) Trypsin/PNGaseF

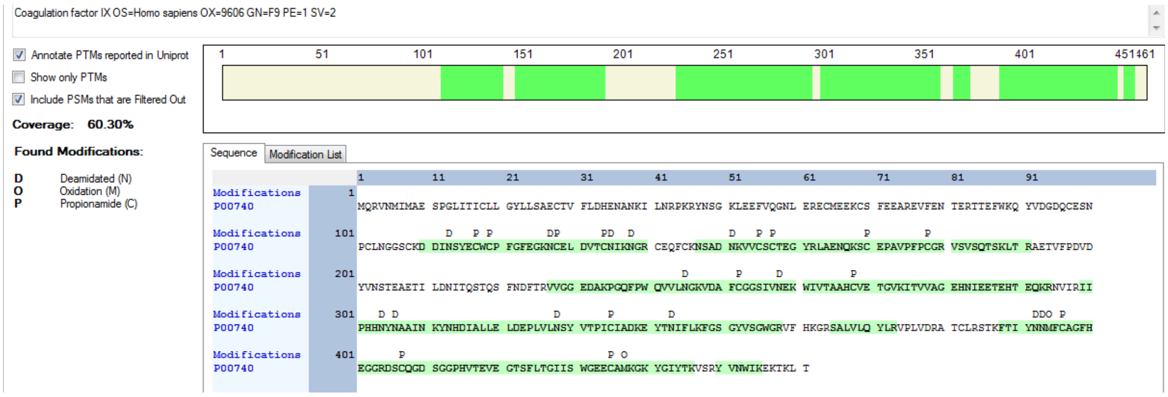


(C) GluC

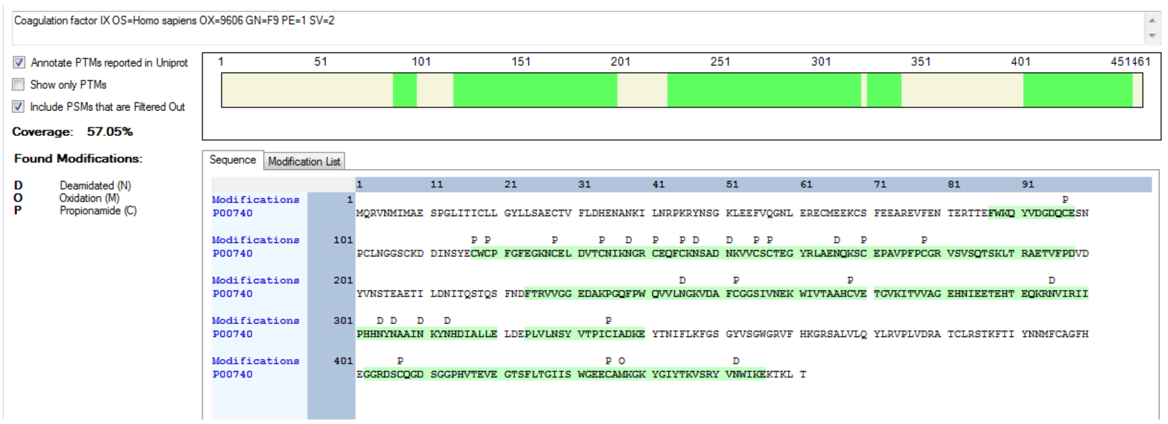

### (D) GluC/PNGase F

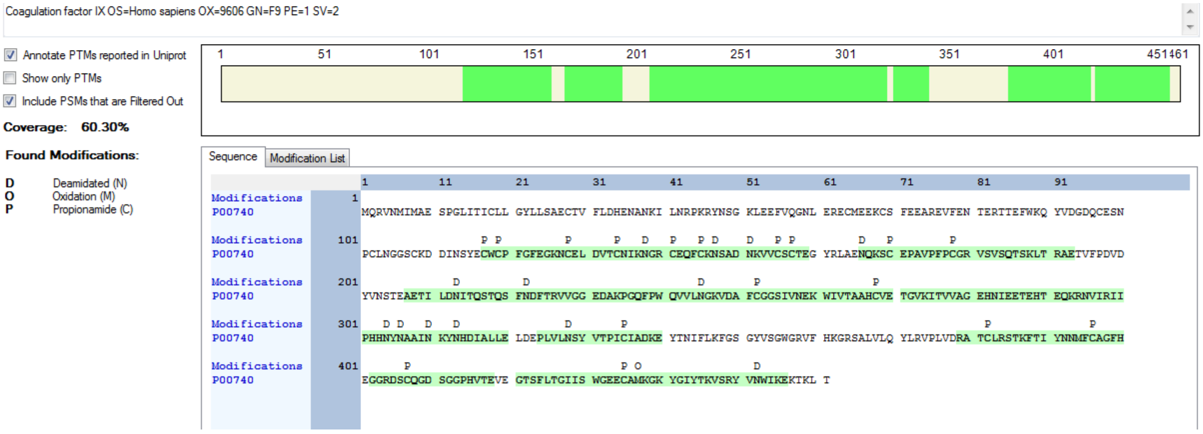


### (E) Chymotrypsin

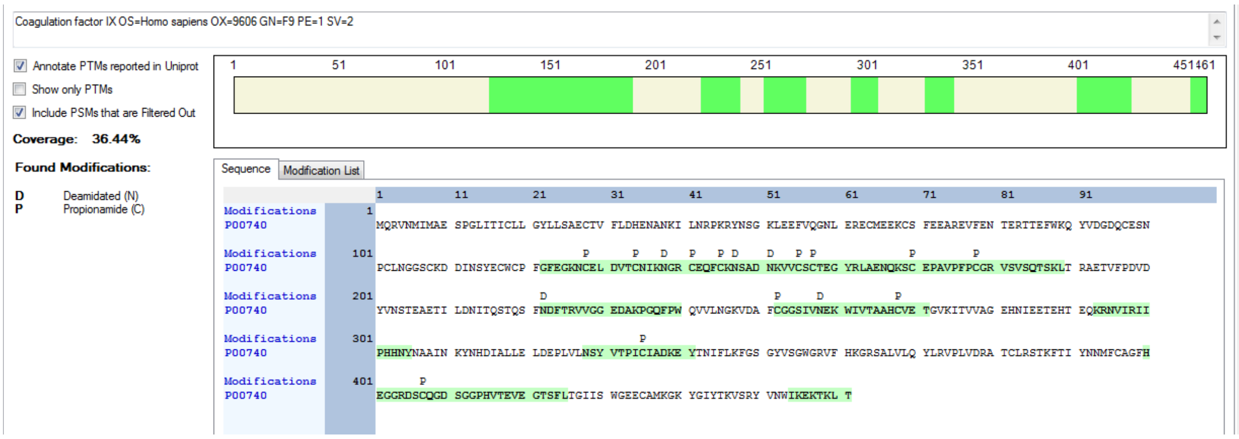


### (F) Chymotrypsin/PNGaseF

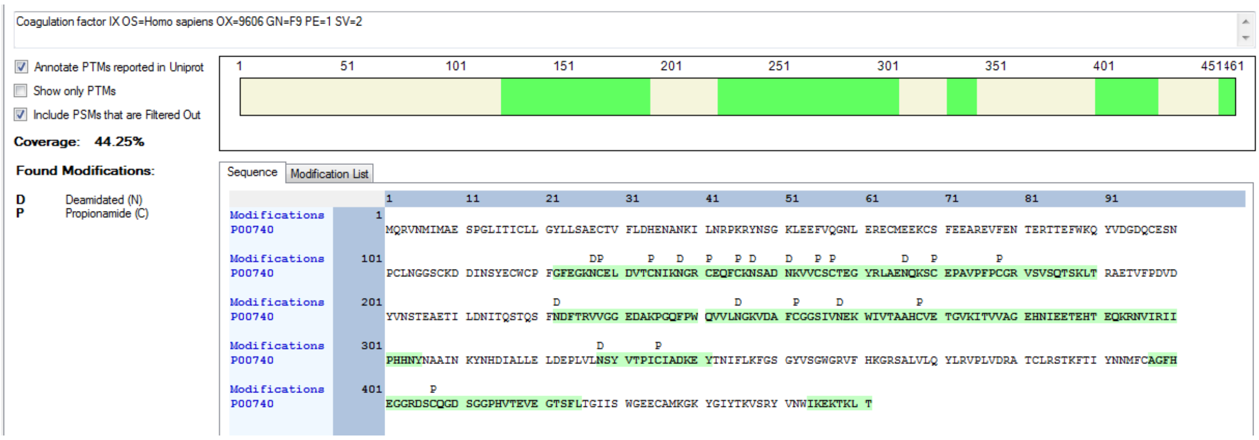

**Figure S-2** Annotated spectra of the novel *O*-linked site of human serum Factor IX from Byonic searches. Spectra have been included from the trypsin, chymotrypsin/PNGase and Glu-C digests of the sample.

Trypsin digested sample:

**(A)** HCD

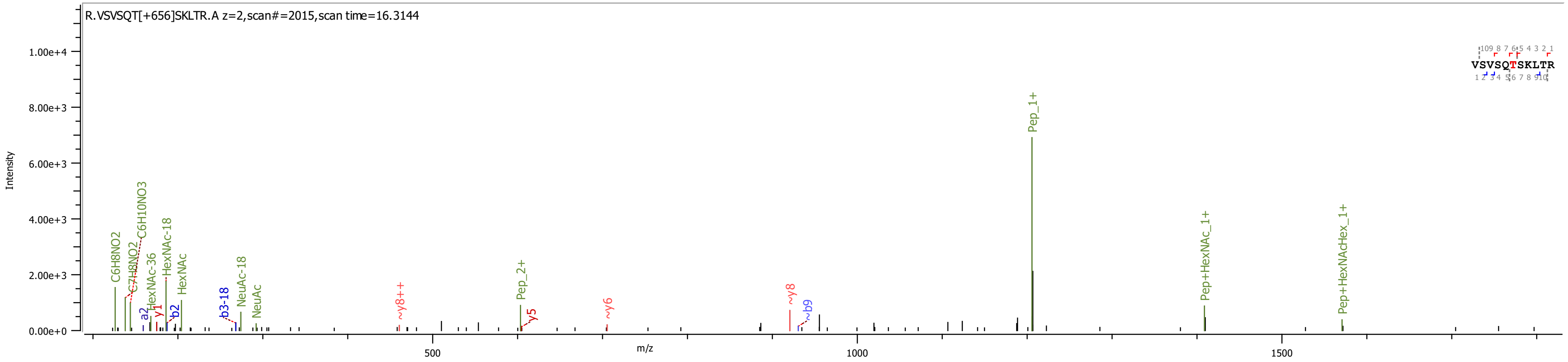


**(B)** HCD-pd_ETciD

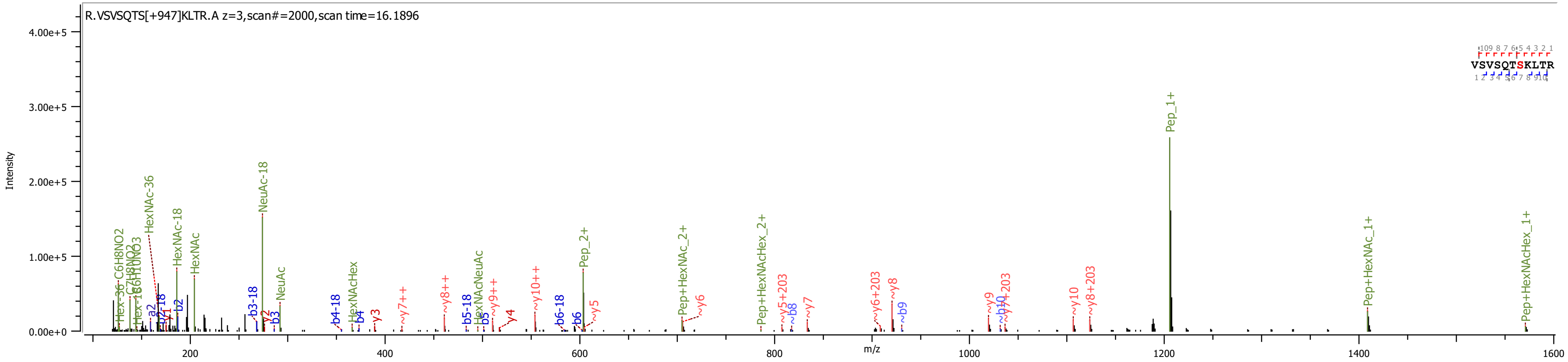

**(C)**

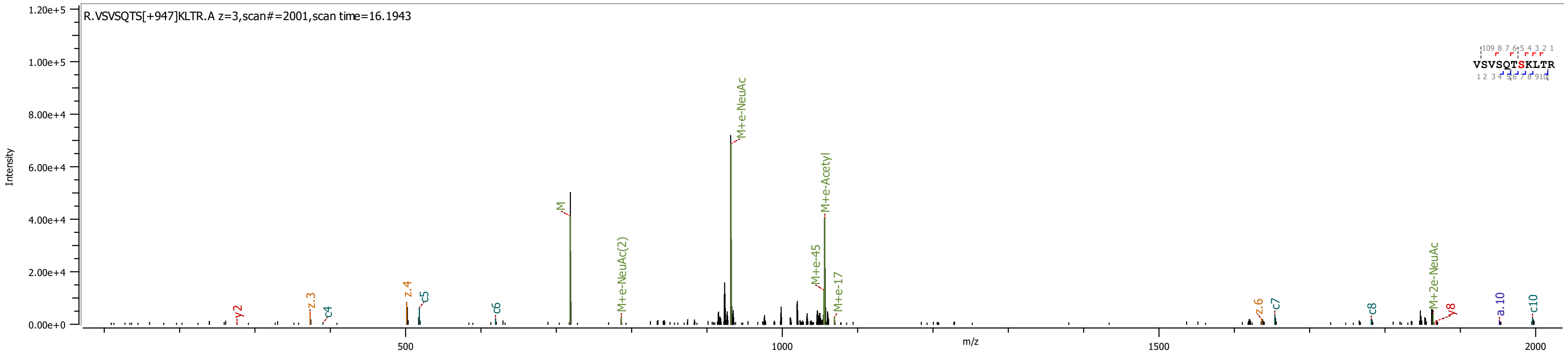


Chymotrypsin/PNGaseF digested sample:

**(D)** HCD-pd-ETciD

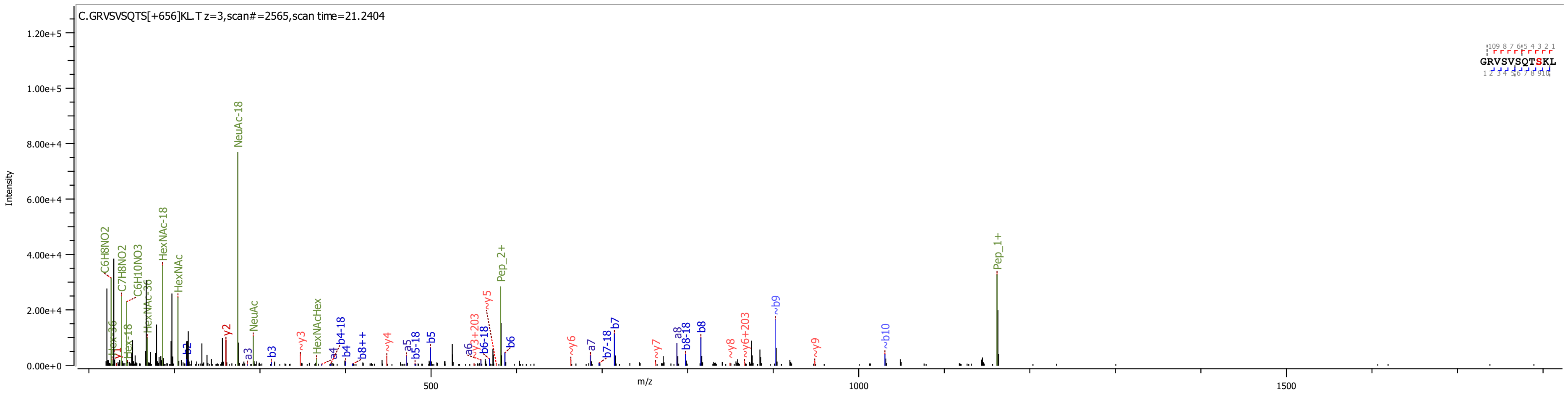

**(E)**

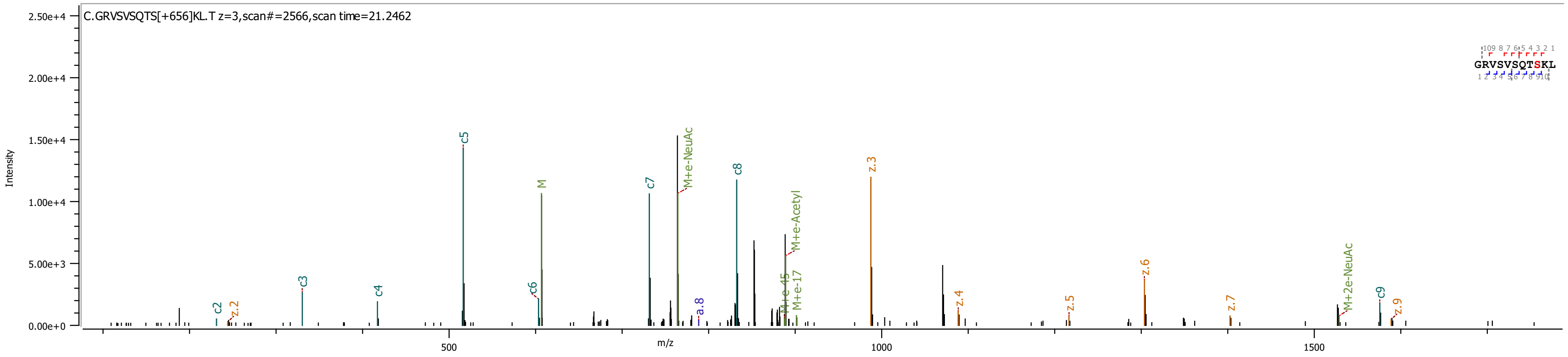


**(F)** HCD-pd-ETciD

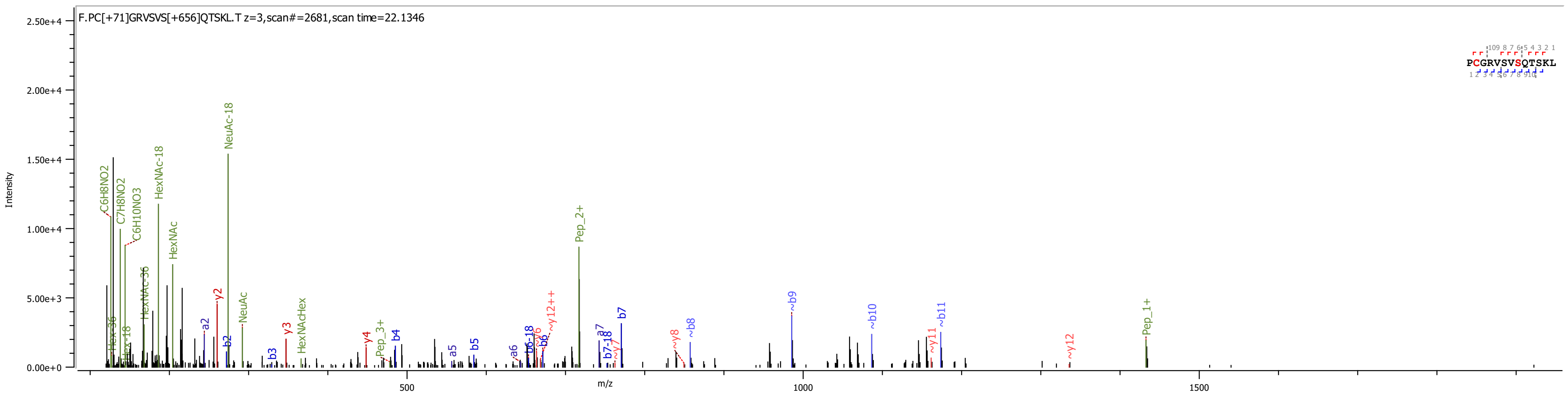

**(G)**

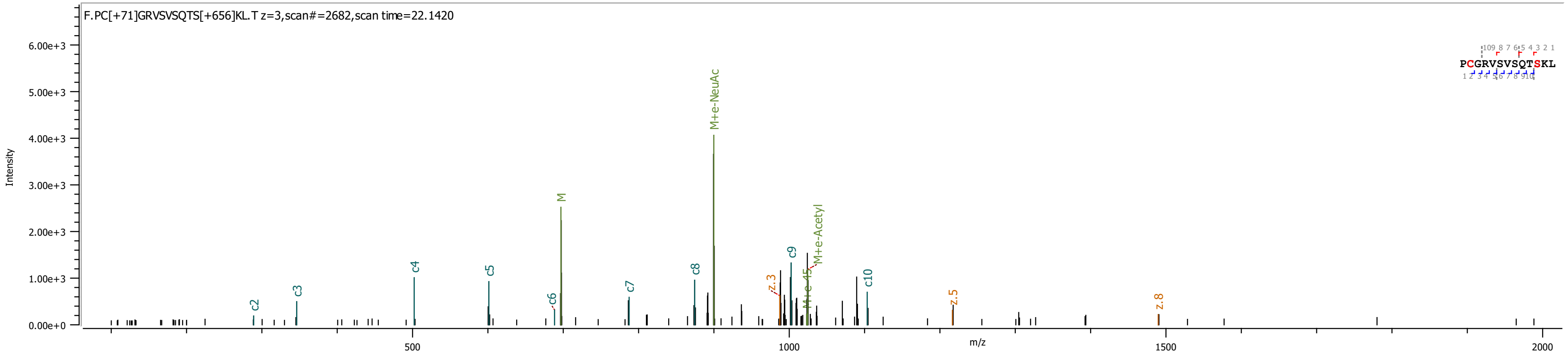


Glu-C digested sample:

**(H)** HCD

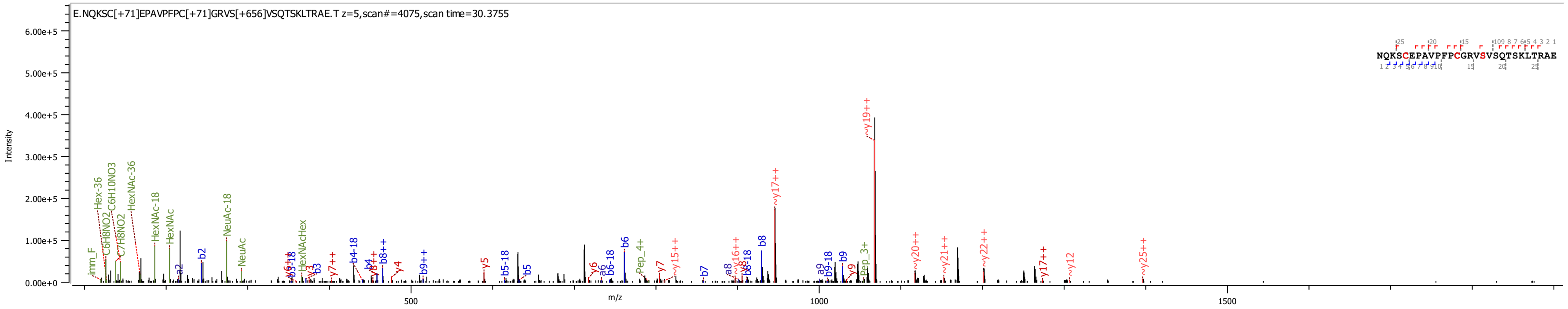

**(I)** HCD

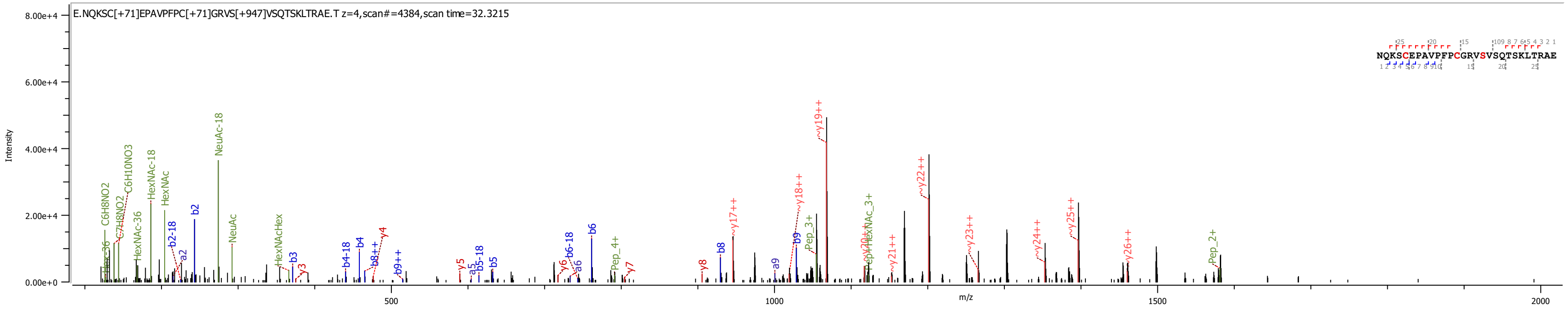


Annotated spectra of the novel *N*-linked site of human serum Factor IX from Byonic searches. Spectra have been included from the trypsin and Glu-C digests of the sample.

Trypsin digested sample:

**(J)** HCD

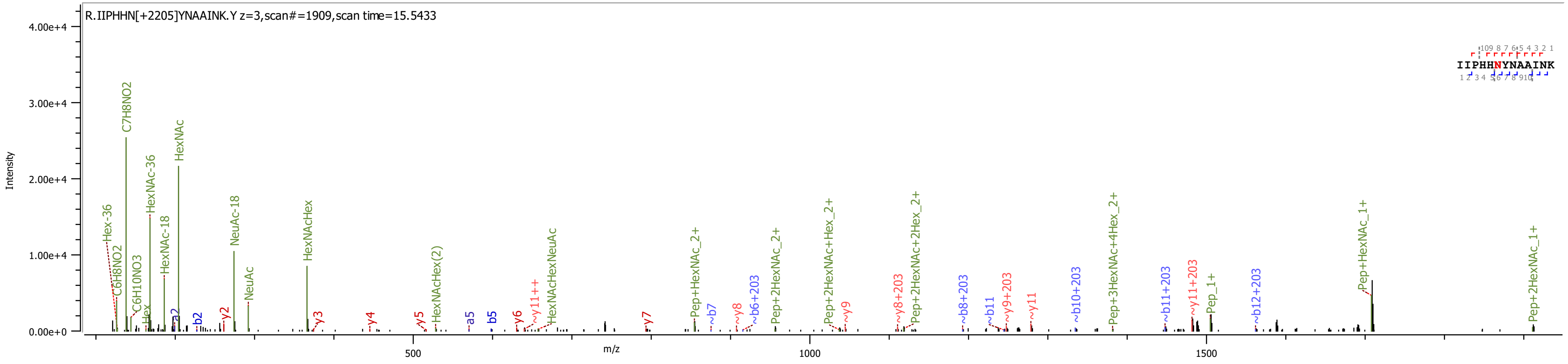

**(K)**

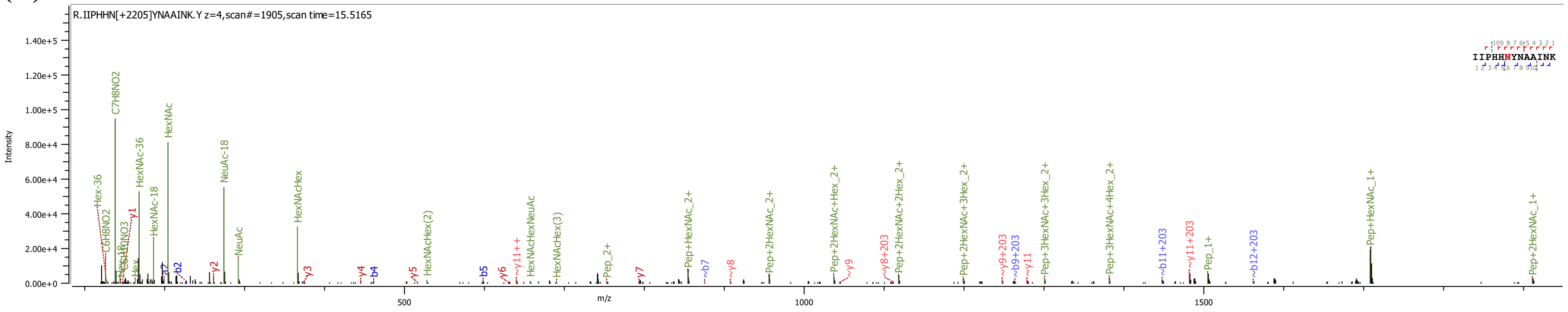


**(L)** HCD-pd_ETciD

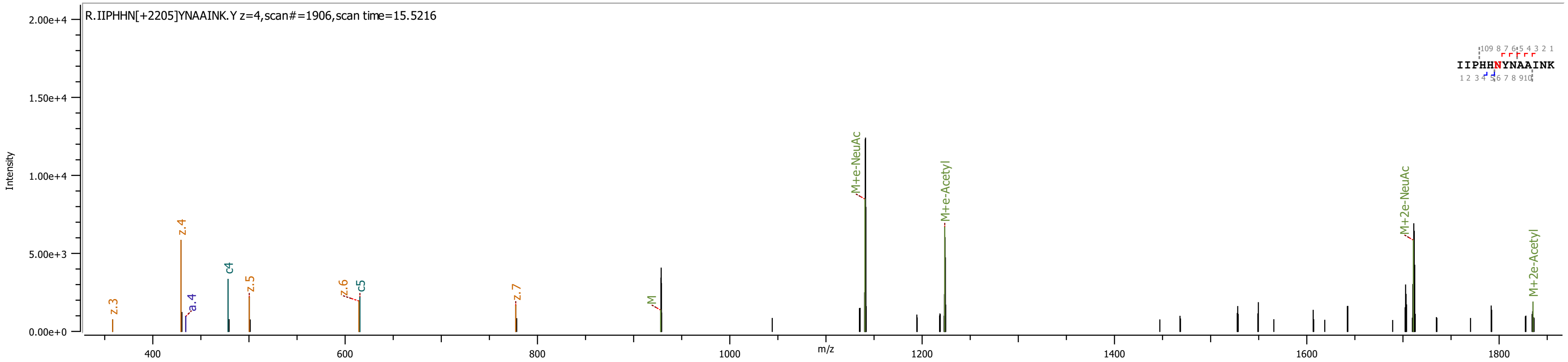

Glu-C digested sample:

**(M)** HCD

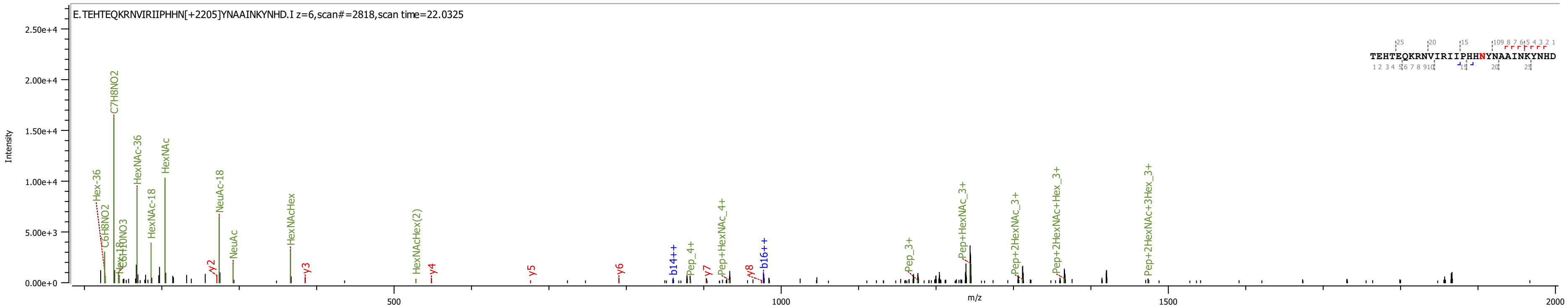

**Figure S-3.** Multiple sequence alignment of human Factor IX, human protein C, human Factor X and human Factor VII and the sequences of Factor IX from Bos taurus (bovine), Pan troglodytes (chimpanzee), Mus musculus (mouse), Canis lupus familiaris (dog), Rattus norvegicus (rat) and Sus scrofa (pig). UniProt accession numbers are listed for each protein and amino acid numbering is based on the full protein sequences.

```
CLUSTAL O(1.2.4) multiple sequence alignment

SP|P00740|FA9_HUMAN  MQRVNMIMAESPGLITICL-L---GYL------------------LSAECTVFLDHENAN 38
SP|P04070|PROC_HUMAN MWQLTS-------LLLFVATWGISGT-------------------PAPLDSVFSSSERAH 34
SP|P00742|FA10_HUMAN MGRPLH-----LVLLSASL-A---GLL------------------L-LGESLFIRREQAN 32
SP|P08709|FA7_HUMAN  ------MVSQ--ALRLLCLLLGLQGCLAAGGVAKASGGETRDMPWKPGPHRVFVTQEEAH 52
SP|P00741|FA9_BOVIN  MWCLNMIMAESPGLVTICL-L---GYL------------------LSAECTVFLDRENAT 38
SP|Q95ND7|FA9_PANTR  MQRVNMIMAESPGLITICL-L---GYL------------------LSAECTVFLDHENAN 38
SP|P16294|FA9_MOUSE  MKHLNTVMAESPALITIFL-L---GYL------------------LSTECAVFLDRENAT 38
SP|P19540|FA9_CANLF  -------MAEASGLVTVCL-L---GYL------------------LSAECAVFLDRENAT 31
SP|P16296|FA9_RAT    -------MADAPGLIPIFL-L---GYL------------------LSTECAVFLDRENAT 31
SP|P16293|FA9_PIG    ------------------------------------------------------------


SP|P00740|FA9_HUMAN  KILNRPKRYNSGKLE-EFVQGNLERECMEEKCSFEEAREVFENTERTTEFWKQYVDGDQC 97
SP|P04070|PROC_HUMAN QVLRIRKRANS--FLEELRHSSLERECIEEICDFEEAKEIFQNVDDTLAFWSKHVDGDQC 92
SP|P00742|FA10_HUMAN NILARVTRANS--FLEEMKKGHLERECMEETCSYEEAREVFEDSDKTNEFWNKYKDGDQC 90
SP|P08709|FA7_HUMAN  GVLHRRRRANA--FLEELRPGSLERECKEEQCSFEEAREIFKDAERTKLFWISYSDGDQC 110
SP|P00741|FA9_BOVIN  KILHRPKRYNSGKLE-EFVRGNLERECKEEKCSFEEAREVFENTEKTTEFWKQYVDGDQC 97
SP|Q95ND7|FA9_PANTR  KILNRPKRYNSGKLE-EFVQGNLERECMEEKCSFEEAREVFENTERTTEFWKQYVDGDQC 97
SP|P16294|FA9_MOUSE  KILTRPKRYNSGKLE-EFVRGNLERECIEERCSFEEAREVFENTEKTTEFWKQYVDGDQC 97
SP|P19540|FA9_CANLF  KILSRPKRYNSGKLE-EFVRGNLERECIEEKCSFEEAREVFENTEKTTEFWKQYVDGDQC 90
SP|P16296|FA9_RAT    KILTRPKRYNSGKLE-EFVQGNLERECIEERCSFEEAREVFENTEKTTEFWKQYVDGDQC 90
SP|P16293|FA9_PIG    --------YNSGKLEESFVRGNLERECIEEKCSFEEAREVFENTEKTNEFWKQYVDGDQC 52
                              *:  :  .:  . ***** ** *.:***:*:*:: : *  ** .: *****

SP|P00740|FA9_HUMAN  ESNP--------CLNGGSCKDDINSYECWCPFGFEGKNCELD----VTCNIKNGRCEQFC 145
SP|P04070|PROC_HUMAN LVLPLEHPCASLCCGHGTCIDGIGSFSCDCRSGWEGRFCQRE-VSFLNCSLDNGGCTHYC 151
SP|P00742|FA10_HUMAN ETSP--------CQNQGKCKDGLGEYTCTCLEGFEGKNCELF--TRKLCSLDNGDCDQFC 140
SP|P08709|FA7_HUMAN  ASSP--------CQNGGSCKDQLQSYICFCLPAFEGRNCETHKDDQLICVNENGGCEQYC 162
SP|P00741|FA9_BOVIN  ESNP--------CLNGGMCKDDINSYECWCQAGFEGTNCELD----ATCSIKNGRCKQFC 145
SP|Q95ND7|FA9_PANTR  ESNP--------CLNGGSCKDDINSYECWCPFGFEGKNCELD----VTCNIKNGRCEQFC 145
SP|P16294|FA9_MOUSE  ESNP--------CLNGGICKDDISSYECWCQVGFEGRNCELD----ATCNIKNGRCKQFC 145
SP|P19540|FA9_CANLF  ESNP--------CLNDGVCKDDINSYECWCRAGFEGKNCELD----VTCNIKNGRCKQFC 138
SP|P16296|FA9_RAT    ESNP--------CLNGGICKDDINSYECWCQAGFEGRNCELD----ATCSIKNGRCKQFC 138
SP|P16293|FA9_PIG    EPNP--------CLNGGLCKDDINSYECWCQVGFEGKNCELD----ATCNIKNGRCKQFC 100
                        *        * . * * * : .: * *  .:**  *:        *  .** * ::*

SP|P00740|FA9_HUMAN  KNSADNKVVCSCTEGYRLAENQKSCEPAVPFPCGRVSVSQTS-KLTRAETVFPD----VD 200
SP|P04070|PROC_HUMAN LEEVG-WRRCSCAPGYKLGDDLLQCHPAVKFPCGRPWKRMEKKRSHLK------------ 198
SP|P00742|FA10_HUMAN HEE-QNSVVCSCARGYTLADNGKACIPTGPYPCGKQTLERRKRSVAQATSSSGEAPDSIT 199
SP|P08709|FA7_HUMAN  SDHTGTKRSCRCHEGYSLLADGVSCTPTVEYPCGKIPILEKRN----------------- 205
SP|P00741|FA9_BOVIN  KRDTDNKVVCSCTDGYRLAEDQKSCEPAVPFPCGRVSVSHISKKLTRAETIFSN----TN 201
SP|Q95ND7|FA9_PANTR  KNSADNKVVCSCTEGYRLAENQKSCEPAVPFPCGRVSVSQTS-KLTRAETVFPD----VD 200
SP|P16294|FA9_MOUSE  KNSPDNKVICSCTEGYQLAEDQKSCEPTVPFPCGRASISYSSKKITRAETVFSN----MD 201
SP|P19540|FA9_CANLF  KLGPDNKVVCSCTTGYQLAEDQRSCEPAVPFPCGRVSVPHISMTRTRAETLFSN----MD 194
SP|P16296|FA9_RAT    KNSPDNKIICSCTEGYQLAEDQKSCEPAVPFPCGRVSVAYNSKKITRAETVFSN----TD 194
SP|P16293|FA9_PIG    KTGADSKVLCSCTTGYRLAPDQKSCKPAVPFPCGRVSVSHSPTTLTRAEIIFSN----MD 156
                              * *  ** *  :   * *:  :***:

SP|P00740|FA9_HUMAN  YVNSTEAET---------ILDNITQSTQSFNDFTRVVGGEDAKPGQFPWQVVL-NGKVDA 250
SP|P04070|PROC_HUMAN ----------------------RDTEDQEDQVDPRLIDGKMTRRGDSPWQVVLLDSKKKL 236
SP|P00742|FA10_HUMAN WKPYDAADLDPTENPFDLLDFNQTQPERGDNNLTRIVGGQECKDGECPWQALLINEENEG 259
SP|P08709|FA7_HUMAN  ----------------------------ASKPQGRIVGGKVCPKGECPWQVLLL-VNGAQ 236
SP|P00741|FA9_BOVIN  YENSSEAEI---------IWDNVTQSNQSFDEFSRVVGGEDAERGQFPWQVLL-HGEIAA 251
SP|Q95ND7|FA9_PANTR  YVNSTEAET---------ILDNITQSTQSFNDFTRVVGGEDAKPGQFPWQVVL-NGKVDA 250
SP|P16294|FA9_MOUSE  YENSTEAVFIQDDITDGAILNNVTESSESLNDFTRVVGGENAKPGQIPWQVIL-NGEIEA 260
SP|P19540|FA9_CANLF  YENSTEVEK---------ILDNVTQ---PLNDFTRVVGGKDAKPGQFPWQVLL-NGKVDA 241
SP|P16296|FA9_RAT    YGNSTEL--ILDDITNSTILDNLTENSEPINDFTRVVGGENAKPGQIPWQVIL-NGEIEA 251
SP|P16293|FA9_PIG    YENSTEVEP---------ILDSLTESNQSSDDFIRIVGGENAKPGQFPWQVLL-NGKIDA 206
                                                   .   *::.*:    *: ***.:*   :

SP|P00740|FA9_HUMAN  FCGGSIVNEKWIVTAAHCVETGVK---ITVVAGEHNIEETEHTEQKRNVIRIIPHHNYNA 307
SP|P04070|PROC_HUMAN ACGAVLIHPSWVLTAAHCMDESKK---LLVRLGEYDLRRWEKWELDLDIKEVFVHPNYS- 292
SP|P00742|FA10_HUMAN FCGGTILSEFYILTAAHCLYQAKR---FKVRVGDRNTEQEEGGEAVHEVEVVIKHNRFT- 315
```

```
SP|P08709|FA7_HUMAN  LCGGTLINTIWVVSAAHCFDKIKNWRNLIAVLGEHDLSEHDGDEQSRRVAQVIIPSTYV- 295
SP|P00741|FA9_BOVIN  FCGGSIVNEKWVVTAAHCIKPGVK---ITVVAGEHNTEKPEPTEQKRNVIRAIPYHSYNA 308
SP|Q95ND7|FA9_PANTR  FCGGSIVNEKWIVTAAHCVDTGVK---ITVVAGEHNIEETEHTEQKRNVIRIIPHHNYNA 307
SP|P16294|FA9_MOUSE  FCGGAIINEKWIVTAAHCLKPGDK---IEVVAGEYNIDKKEDTEQRRNVIRTIPHHQYNA 317
SP|P19540|FA9_CANLF  FCGGSIINEKWVVTAAHCIEPDVK---ITIVAGEHNTEKREHTEQKRNVIRTILHHSYNA 298
SP|P16296|FA9_RAT    FCGGAIINEKWIVTAAHCLKPGDK---IEVVAGEHNIDEKEDTEQRRNVIRTIPHHQYNA 308
SP|P16293|FA9_PIG    FCGGSIINEKWVVTAAHCIEPGVK---ITVVAGEYNTEETEPTEQRRNVIRAIPHHSYNA 263
                      **. ::   ::::****.    .   :    *: :  . :  *    :   :    :

SP|P00740|FA9_HUMAN  AINKYNHDIALLELDEPLVLNSYVTPICIADKEYTNI-FLKFG-SGYVSGWGRVFHKGR- 364
SP|P04070|PROC_HUMAN -KSTTDNDIALLHLAQPATLSQTIVPICLPDSGLAERELNQAGQETLVTGWGYHSSREKE 351
SP|P00742|FA10_HUMAN -KETYDFDIAVLRLKTPITFRMNVAPACLPERDWAESTLMTQK-TGIVSGFGRTHEKGR- 372
SP|P08709|FA7_HUMAN  -PGTTNHDIALLRLHQPVVLTDHVVPLCLPERTFSERTLAFVR-FSLVSGWGQLLDRGA- 352
SP|P00741|FA9_BOVIN  SINKYSHDIALLELDEPLELNSYVTPICIADRDYTNI-FLKFG-YGYVSGWGKVFNRGR- 365
SP|Q95ND7|FA9_PANTR  AINKYNHDIALLELDEPLVLNSYVTPICIADKEYTNI-FLKFG-SGYVSGWGRVFHKGR- 364
SP|P16294|FA9_MOUSE  TINKYSHDIALLELDKPLILNSYVTPICVANREYTNI-FLKFG-SGYVSGWGKVFNKGR- 374
SP|P19540|FA9_CANLF  TINKYNHDIALLELDEPLTLNSYVTPICIADREYSNI-FLKFG-SGYVSGWGRVFNKGR- 355
SP|P16296|FA9_RAT    TINKYSHDIALLELDKPLILNSYVTPICVANKEYTNI-FLKFG-SGYVSGWGKVFNKGR- 365
SP|P16293|FA9_PIG    TVNKYSHDIALLELDEPLTLNSYVTPICIADKEYTNI-FLKFG-SGYVSGWGRVFNRGR- 320
                        . . ***:*.*  *  :   :.* *: :   ::  :        *:*:*    :

SP|P00740|FA9_HUMAN  ----SALVLQYLRVPLVDRATCLRST-----KFTIYNNMFCAGFHEGGRDSCQGDSGGPH 415
SP|P04070|PROC_HUMAN AKRNRTFVLNFIKIPVVPHNECSEVM-----SNMVSENMLCAGILGDRQDACEGDSGGPM 406
SP|P00742|FA10_HUMAN ----QSTRLKMLEVPYVDRNSCKLSS-----SFIITQNMFCAGYDTKQEDACQGDSGGPH 423
SP|P08709|FA7_HUMAN  ----TALELMVLNVPRLMTQDCLQQSRKVGDSPNITEYMFCAGYSDGSKDSCKGDSGGPH 408
SP|P00741|FA9_BOVIN  ----SASILQYLKVPLVDRATCLRST-----KFSIYSHMFCAGYHEGGKDSCQGDSGGPH 416
SP|Q95ND7|FA9_PANTR  ----SALVLQYLRVPLVDRATCLRST-----KFTIYNNMFCAGFHEGGRDSCQGDSGGPH 415
SP|P16294|FA9_MOUSE  ----QASILQYLRVPLVDRATCLRST-----TFTIYNNMFCAGYREGGKDSCEGDSGGPH 425
SP|P19540|FA9_CANLF  ----SASILQYLKVPLVDRATCLRST-----KFTIYNNMFCAGFHEGGKDSCQGDSGGPH 406
SP|P16296|FA9_RAT    ----QASILQYLRVPLVDRATCLRST-----KFSIYNNMFCAGYREGGKDSCEGDSGGPH 416
SP|P16293|FA9_PIG    ----SATILQYLKVPLVDRATCLRST-----KVTIYSNMFCAGFHEGGKDSCLGDSGGPH 371
                          :  *  :.:* :    *         .  : . *:***     .*:* ******

SP|P00740|FA9_HUMAN  VTEVEGTSFLTGIISWGEECAMKGKYGIYTKVSRYVNWIKEKTKLT-------------- 461
SP|P04070|PROC_HUMAN VASFHGTWFLVGLVSWGEGCGLLHNYGVYTKVSRYLDWIHGHIRDKEAPQKSW-AP---- 461
SP|P00742|FA10_HUMAN VTRFKDTYFVTGIVSWGEGCARKGKYGIYTKVTAFLKWIDRSMKTRGLPKAKSHAPEVIT 483
SP|P08709|FA7_HUMAN  ATHYRGTWYLTGIVSWGQGCATVGHFGVYTRVSQYIEWLQKLMRSEPRPGVLLRAPFP-- 466
SP|P00741|FA9_BOVIN  VTEVEGTSFLTGIISWGEECAMKGKYGIYTKVSRYVNWIKEKTKLT-------------- 462
SP|Q95ND7|FA9_PANTR  VTEVEGTSFLTGIISWGEECAMKGKYGIYTKVSRYVNWIKEKTKLT-------------- 461
SP|P16294|FA9_MOUSE  VTEVEGTSFLTGIISWGEECAMKGKYGIYTKVSRYVNWIKEKTKLT-------------- 471
SP|P19540|FA9_CANLF  VTEVEGISFLTGIISWGEECAMKGKYGIYTKVSRYVNWIKEKTKLT-------------- 452
SP|P16296|FA9_RAT    VTEVEGTSFLTGIISWGEECAMKGKYGIYTKVSRYVNWIKEKTKLT-------------- 462
SP|P16293|FA9_PIG    VTEVEGTSFLTGIISWGEECAVKGKYGIYTKVSRYVNW---------------------- 409
                     .:  ..  ::.*::***: *.   ::*:**:*: ::.*

SP|P00740|FA9_HUMAN  -----
SP|P04070|PROC_HUMAN -----
SP|P00742|FA10_HUMAN SSPLK 488
SP|P08709|FA7_HUMAN  -----
SP|P00741|FA9_BOVIN  -----
SP|Q95ND7|FA9_PANTR  -----
SP|P16294|FA9_MOUSE  -----
SP|P19540|FA9_CANLF  -----
SP|P16296|FA9_RAT    -----
SP|P16293|FA9_PIG    -----
```

**Fig. S-4** Crystal structure of human Factor IXa. Surface representation of Factor IXa with cartoon representation of the protein backbone (Homology modelled from PDB 1CF1, 1EDNM and 3KCG and taken from http://www.factorix.org/structure.html.php). The colouring of the domains follows that of Fig. 1 within the manuscript. The GLA domain (red) anchors onto phospholipid membranes and the EGF1 (pink) and EGF2 (purple) domains form a stalk. The activation peptide is not present in this structure. The serine protease domain (orange) contains the active site (yellow spheres) and a surface loop (magenta) within the protease domain includes an Asn residue (magenta spheres) within a highly conserved *N*-glycosylation sequon.

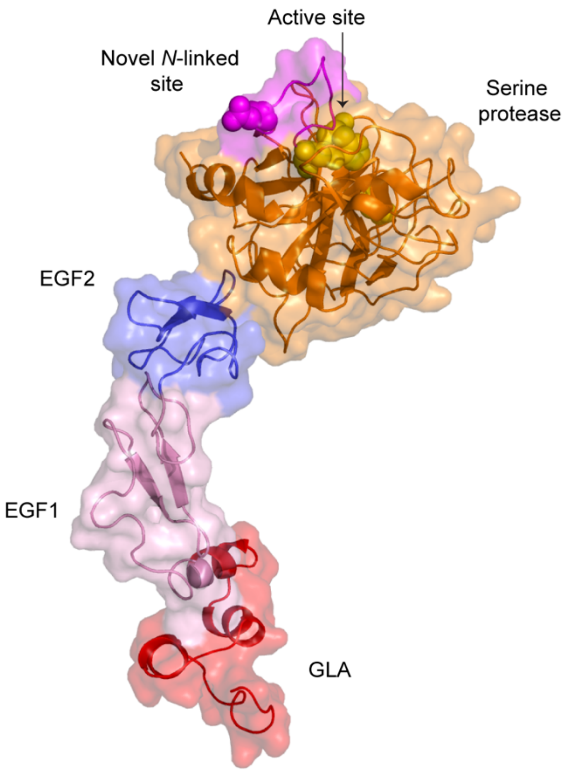

**Figure S-5** Annotated spectra of reported post-translational modifications (PTMs) of Factor IX from Byonic searches.

(A) Reported modification of γ-carboxylation at E40[1-2] from the trypsin digested sample:

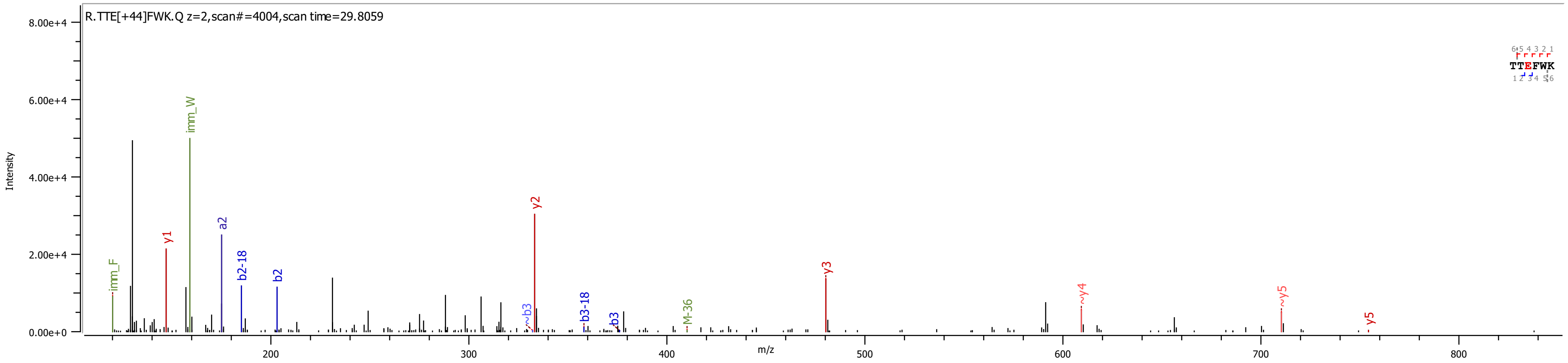


(B) Reported modification of β-hydroxylation of Asp64[3-4] from the trypsin digested sample.

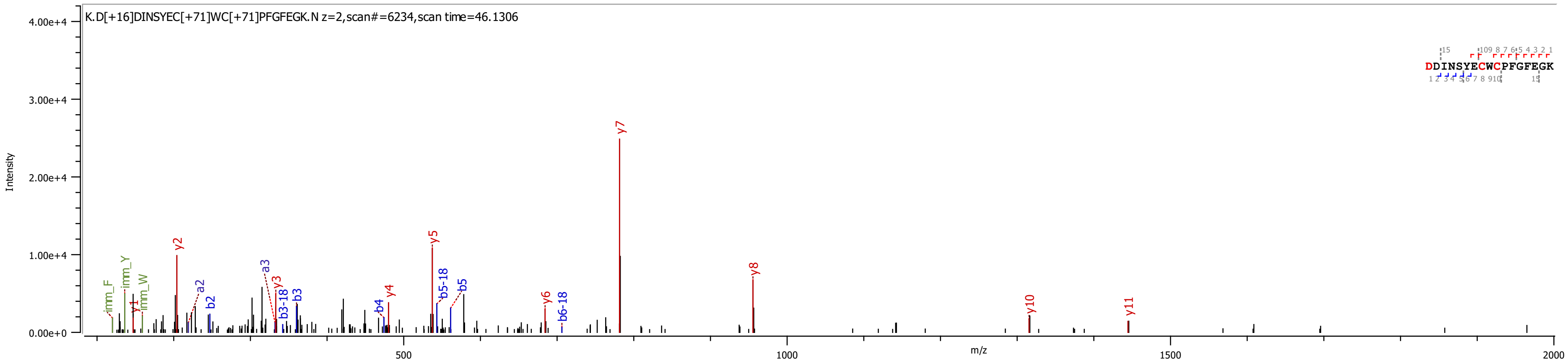

(C) Reported modifications of *O*-linked glucose at Ser53[5], *O*-linked fucose at Ser61[6-7] and β-hydroxylation of Asp64[3-4] from the Glu-C digested sample. Please note Byonic has assigned the glycans in different positions to those reported in the literature.

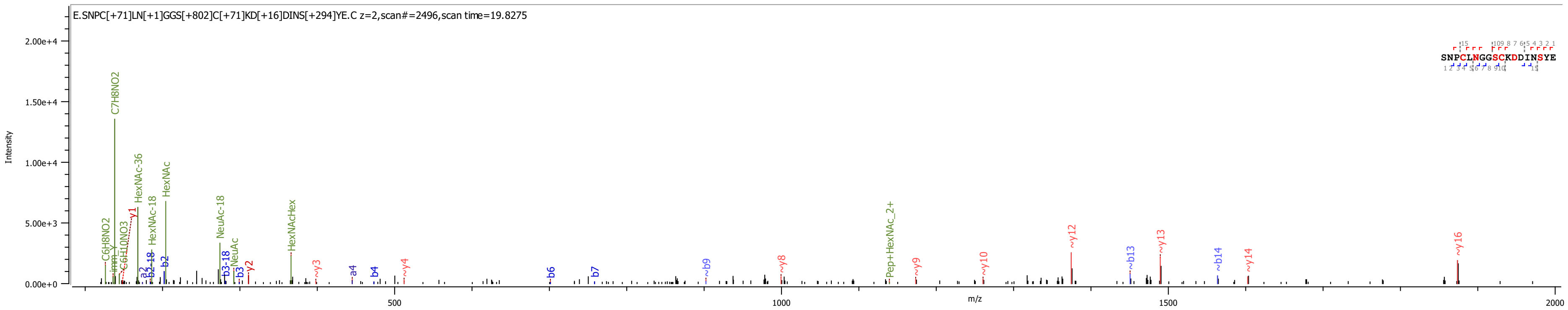


(D) Reported modifications of *O*-linked glucose at Ser53[5] and *O*-linked fucose at Ser61[6-7] from the Glu-C digested sample.

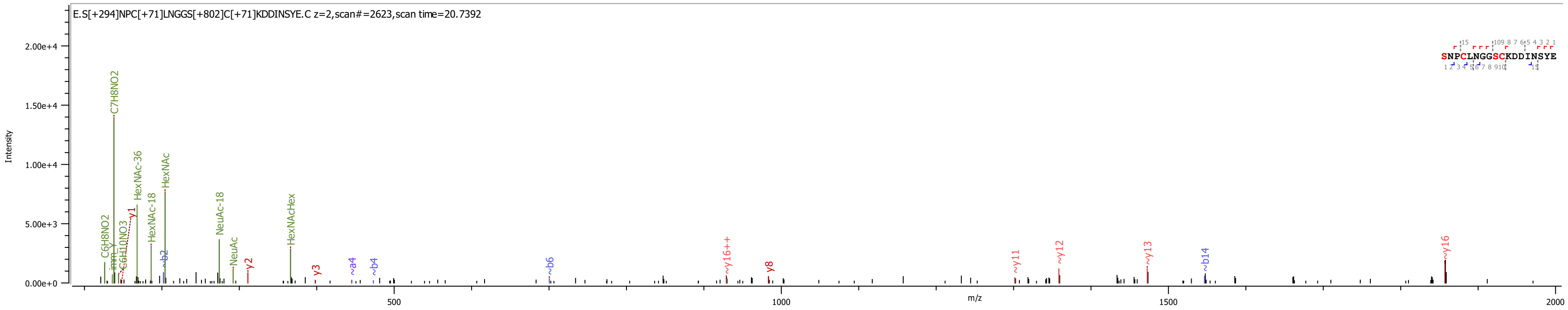

(E) Reported modification of *O*-linked glucose at Ser61[6-7] from the chymotrypsin/PNGaseF digested sample.

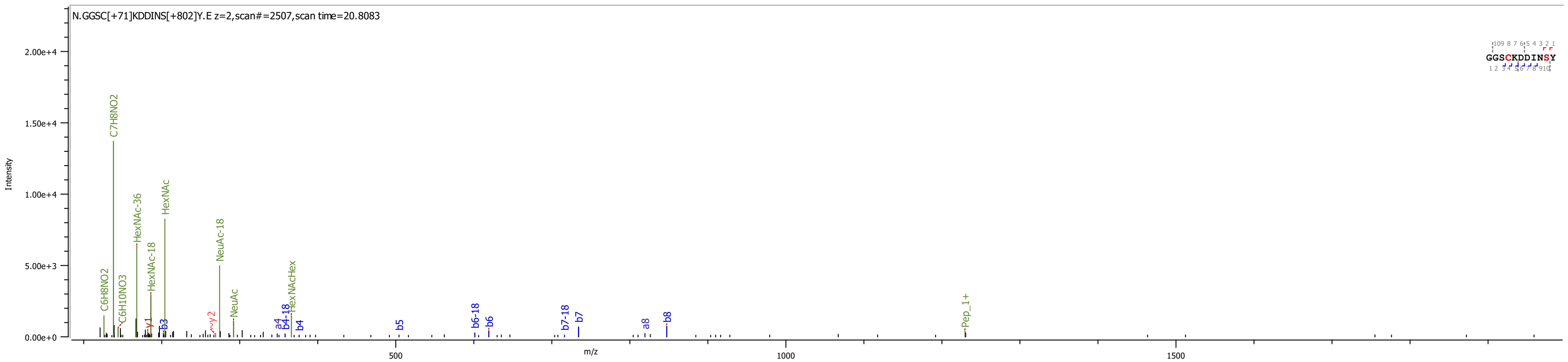


(F) Reported modification of phosphorylation at S68 (Noted in UniProt identifier P00740) from the trypsin digested sample.

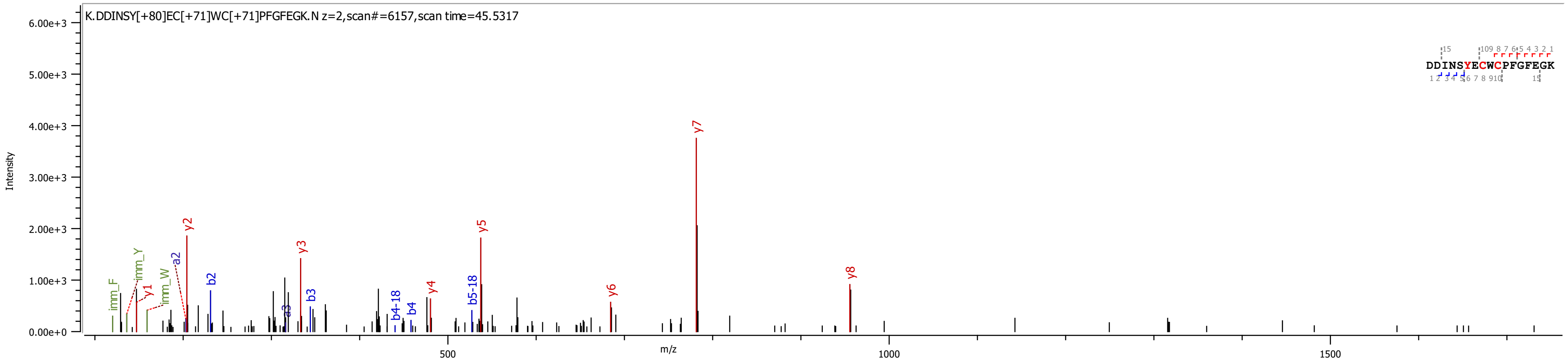

### References


1. Stanley, T. B.; Wu, S.-M.; Houben, R. J. T. J.; Mutucumarana, V. P.; Stafford, D. W., Role of the propeptide and γ-glutamic acid domain of factor ix for in vitro carboxylation by the vitamin k-dependent carboxylase. *Biochemistry* **1998,** *37* (38), 13262-13268.
2. Jorgensen, M. J.; Cantor, A. B.; Furie, B. C.; Brown, C. L.; Shoemaker, C. B.; Furie, B., Recognition site directing vitamin K-dependent gamma-carboxylation resides on the propeptide of factor IX. *Cell* **1987,** *48* (2), 185-91.
3. Derian, C. K.; VanDusen, W.; Przysiecki, C. T.; Walsh, P. N.; Berkner, K. L.; Kaufman, R. J.; Friedman, P. A., Inhibitors of 2-ketoglutarate-dependent dioxygenases block aspartyl beta-hydroxylation of recombinant human factor IX in several mammalian expression systems. *J. Biol. Chem.* **1989,** *264* (12), 6615-18.
4. Fernlund, P.; Stenflo, J., Beta-hydroxyaspartic acid in vitamin K-dependent proteins. *J. Biol. Chem.* **1983,** *258* (20), 12509-12.
5. Nishimura, H.; Kawabata, S.; Kisiel, W.; Hase, S.; Ikenaka, T.; Takao, T.; Shimonishi, Y.; Iwanaga, S., Identification of a disaccharide (Xyl-Glc) and a trisaccharide (Xyl2-Glc) O-glycosidically linked to a serine residue in the first epidermal growth factor-like domain of human factors VII and IX and protein Z and bovine protein Z. *J. Biol. Chem.* **1989,** *264* (34), 20320-5.
6. Harris, R. J.; van Halbeek, H.; Glushka, J.; Basa, L. J.; Ling, V. T.; Smith, K. J.; Spellman, M. W., Identification and structural analysis of the tetrasaccharide NeuAcα (2→ 6) Galβ (1→ 4) GlcNAcβ (1→ 3) Fucα1→ O-linked to serine 61 of human factor IX. *Biochemistry* **1993,** *32* (26), 6539-47.
7. Nishimura, H.; Takao, T.; Hase, S.; Shimonishi, Y.; Iwanaga, S., Human factor IX has a tetrasaccharide O-glycosidically linked to serine 61 through the fucose residue. *J. Biol. Chem.* **1992,** *267* (25), 17520-25.
